# KDM2B controls HIF levels and activity through its JmjC and CxxC domains

**DOI:** 10.64898/2026.03.26.714448

**Authors:** Michael Batie, Dilem Shakir, Chun-Sui Kwok, Gemma Bell, Jiahua Kou, Ali Bakhsh, Sonia Rocha

**Affiliations:** Department of Biochemistry, Cell and Systems Biology, Institute of Systems, Molecular and Integrative Biology, University of Liverpool., Biosciences Building, Crown Street, Liverpool L697ZB, United Kingdom

**Author notes:** Corresponding authors: Michael Batie, Sonia Rocha.

## Abstract

Hypoxia-inducible factors (HIFs) are key regulators of cellular responses to low oxygen (hypoxia), controlling the expression of genes required for survival and adaptation. KDM2B, a chromatin-modifying enzyme, is a direct target of HIF-1α, but its precise role in regulating HIF and the hypoxia response remains unclear. Here, we investigated the role of KDM2B in the response to hypoxia in a variety of cell lines. Our analysis reveals that KDM2B depletion regulates HIF activity in a cell type-dependent manner, with KDM2B depletion decreasing HIF activity in U2OS and MDA-MB-231 cells and increasing HIF activity in HeLa cells. We show that KDM2B depletion also reduces HIF-1α protein and RNA expression and reduces HIF-1α binding at hypoxia-response elements of its target genes in U2OS and MDA-MB-231 cells. Conversely, overexpression of KDM2B enhances HIF activity and HIF-1α levels in both U2OS and HEK293 cells. Mechanistically, we find that KDM2B requires its JmjC demethylase and CxxC DNA-binding domains for HIF regulation. Furthermore, we demonstrate that KDM2B is required for RNA Pol II recruitment to the promoter of *HIF1A*. At the cellular level, KDM2B supports cell proliferation, with its depletion impairing proliferation and reducing cell numbers under hypoxic conditions. Our work highlights a new function of KDM2B as a key regulator of HIF-1α expression, acting through its demethylase and DNA-binding functions. Our data indicate that KDM2B is essential for cellular adaptation to hypoxia, impacting both HIF-dependent gene expression and cell survival, and has important implications for our understanding of HIF regulation.

## Introduction

The ability to sense, respond, and adapt to changes in molecular oxygen availability is fundamental to all metazoans [1]. Reduced oxygen availability (hypoxia) induces a variety of adaptive cellular responses, which are orchestrated in part by coordinated changes in transcriptional programmes [2]. Hypoxia Inducible Factors (HIFs) are critical regulators of transcriptional response to hypoxia. The HIF family contains oxygen-sensitive α subunits (HIF-1α, HIF-2α, HIF-3α) and a constitutively expressed β subunit (HIF-1β), and they typically function as HIF-1β containing heterodimers [3]. HIFs are directly regulated by oxygen through the class of 2-OG dependent dioxygenases (2-OGDDs) called Prolyl Hydroxylases (PHD1-3) [4]. Under normal oxygen tensions, PHDs hydroxylate key proline residues within the oxygen-dependent degradation domain of HIF-1α subunits, leading to enhanced affinity for the Von Hippel Lindau (VHL) containing RBX/Cullin2 E3 ligase, which catalyses Lysine-48 ubiquitination and proteasomal degradation of HIF-1α subunits [5]. PHDs are cellular oxygen sensors, and their activity is impaired in hypoxia, due to their dependence and sensitivity to molecular oxygen; this results in HIF-α stabilisation and activation of the HIF pathway [5, 6]. HIFs bind hypoxia response elements (HREs) and control the expression of genes involved in a range of cellular processes, including metabolism, angiogenesis, autophagy, and cell cycle control [2, 3, 7]. In addition to protein stability, HIFs have also been shown to be modulated by various mechanisms at the transcriptional and translational level [8–10]. For example, HIF transcription is controlled by the NF-κB family of transcription factors. This occurs in response to cytokines [11–14], infection [13], and reactive oxygen species [15].

Jumonji-C (JmjC) domain containing proteins are a large subfamily of 2-OGDDs that require oxygen for their enzymatic activity [16, 17]. JmjC domain containing proteins are functionally important in mediating the hypoxia response. Inactivation of KDM5A and KDM6A in hypoxia is critical for hypoxia-stimulated processes, including cell proliferation and differentiation of stem cells, respectively [18, 19]. Furthermore, several JmjC domain containing proteins are hypoxia-inducible HIF target genes, displaying increased expression in hypoxia [5, 20, 21]. Interestingly, some of these hypoxia-induced JmjC domain containing proteins, namely KDM3A and KDM4C, remain active at low oxygen concentrations and function in hypoxia as HIF coactivators through their histone demethylase activity [22–24].

We have previously demonstrated that the KDM2 subfamily of JmjC domain containing proteins (KDM2A and KDM2B) are HIF-1 target genes in different human cancer cells [25]. However, whether the KDM2 family members contribute to the cellular response to hypoxia is currently unknown.

KDM2B is a multi-functional chromatin-binding protein with critical roles in stem cell biology and development and is highly dysregulated across various cancer types [16, 26–28]. Here, we investigated the role of KDM2B in modulating HIF and the hypoxia response in several human cell lines. Utilising HIF activity luciferase reporter assays, qPCR, and Western blotting, we find that KDM2B can function as a positive regulator of HIF-1α and is required for full activation of HIF-1α levels and activity in response to hypoxia, in some of the cell lines studied. Furthermore, we show that overexpression of KDM2B can increase HIF-1α levels and HIF activity. Mechanistically, we demonstrate that KDM2B regulates HIF-1α through its JmjC and CxxC domains and controls HIF-1α RNA expression and RNA Pol II recruitment to the *HIF1A* promoter. Interestingly, we also observe that KDM2B positively regulates NF-κB-dependent reporter activity in response to TNF-α. Finally, we find that KDM2B is required for the proliferation of cells in normoxia and hypoxia.

## Results

### KDM2B regulates HIF activity in a cell type dependent manner

HIFs are central regulators of the hypoxia response [3]. We previously identified the KDM2 family (KDM2A and KDM2B) of JmjC domain containing proteins as hypoxia-upregulated HIF-1 target genes [25]. Other hypoxia-upregulated JmjC domain containing proteins can function as HIF coactivators [22–24]. However, the regulation of HIF and the hypoxia response by KDM2s has not yet been determined. To initially investigate potential HIF regulation by KDM2B, we depleted KDM2B using 2 different siRNAs in three human cancer cell lines (U2OS, MDA-MB-231, and HeLa) stably expressing a HIF-dependent luciferase reporter (HRE-luciferase) (Figure 1A), exposed the cells or not to 24 hours of 1% oxygen as a hypoxia treatment, and measured HIF activity by luciferase assays (Figure 1B). We chose these specific cell lines as they have well-characterised responses to hypoxia [29] and we have previously demonstrated that hypoxia induces KDM2B expression in U2OS and HeLa cells [25]. Control oxygen conditions were normal cell culture levels of 21%, as these cells have been acclimatised to sea level oxygen since being established. As expected, HIF-dependent luciferase activity increased in control siRNA-treated cells from all cell lines in response to hypoxia (Figure 1B). Depletion of KDM2B reduced HIF-dependent luciferase activity in response to hypoxia in U2OS and MDA-MB-231 HRE-luciferase cells (Figure 1B). Interestingly, in HeLa HRE-luciferase cells, depletion of KDM2B resulted in increased HIF-dependent luciferase activity (Figure 1B), demonstrating that regulation of HIF by KDM2B is to some extent cell type dependent. Due to difficulties in detecting endogenous protein levels of KDM2B in these cell lines using Western blotting [25], we validated siRNA-mediated depletion of KDM2B in each of the cell lines by qPCR (Figure 1C). To understand if this reporter activity regulation is dynamic, we performed gain-of-function experiments in U2OS and HeLa HRE-luciferase cells. In both cell lines, overexpression of KDM2B led to an increase in HIF-dependent luciferase activity under normal oxygen levels (21% oxygen) (Figure 1D). Taken together, these results indicate that KDM2B can modulate HIF activity in a cell type dependent manner.

**Figure 1.**
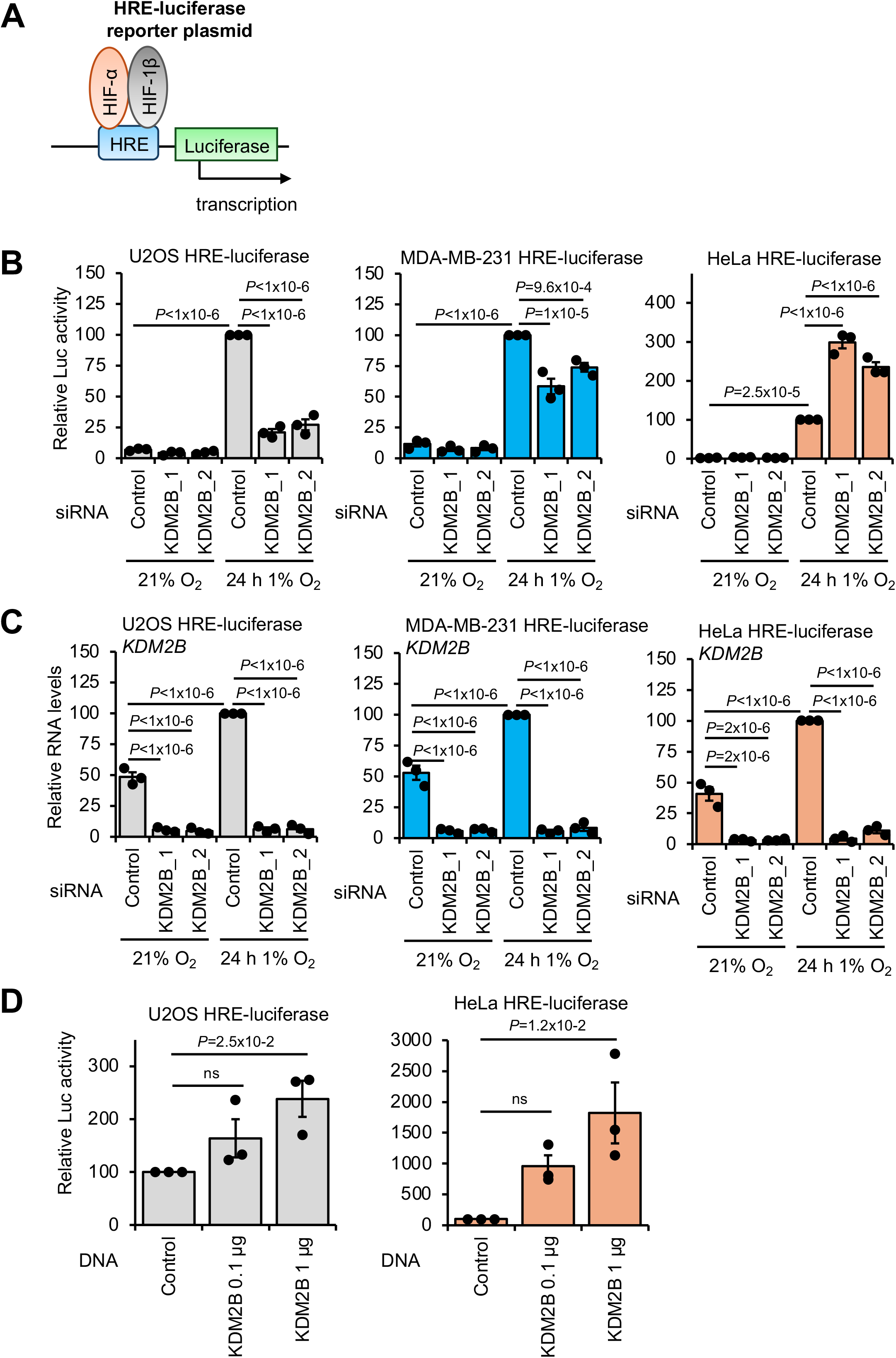
KDM2B is a positive regulator of HIF reporter activity. **A)** Schematic of the hypoxia response element (HRE)-luciferase plasmid stably expressed in cells to measure HIF activity. **B)** Luciferase assay analysis in U2OS, MDA-MB-231, and HeLa HRE-luciferase cells cultured at 21% oxygen, transfected with control, KDM2B_1, or KDM2B_2 siRNA for 48 hours, and exposed or not to 1% oxygen for 24 hours. Data represent mean (n=3 biological replicates) ± SEM. *P* values for group comparisons were calculated by one-way ANOVA with Tukey’s test. **C)** qPCR analysis of *KDM2B* RNA levels in U2OS, MDA-MB-231, and HeLa HRE-luciferase cells cultured at 21% oxygen, transfected with control, KDM2B_1, or KDM2B_2 siRNA for 48 hours, and exposed or not to 1% oxygen for 24 hours. *ACTB* was used as a normalising gene. Data represent mean (n=3 biological replicates) ± SEM. *P* values for group comparisons were calculated by one-way ANOVA with Tukey’s test. **D)** Luciferase assay analysis in U2OS and HeLa HRE-luciferase cells cultured at 21% oxygen and transfected with control, KDM2B (0.1 µg), or KDM2B (1 µg) plasmids for 48 hours. Data represent mean (n=3 biological replicates) ± SEM. *P* values for group comparisons were calculated by one-way ANOVA with Dunnett’s test.

Given that luciferase reporter analysis, although useful, is artificial in nature, we next investigated the effect of KDM2B depletion on the protein expression of HIF target genes by Western blotting (Figure 2A, Supplementary Figure S1-S2). As HeLa cells have the same response to depletion and overexpression of KDM2B, we performed the majority of our subsequent analyses on the other cell types. As expected, protein levels of the core HIF target genes, CAIX (gene name *CA9*), BNIP3L, and BNIP3, were upregulated in control siRNA-treated U2OS and MDA-MB-231 cells exposed to hypoxia (24 hours 1% oxygen) (Figure 2A, Supplementary Figure S1). In both MDA-MB-231 and U2OS cells, loss of KDM2B reduced the protein levels of CAIX, BNIP3L, and BNIP3 in hypoxia (Figure 2A, Supplementary Figure S1), supporting the HIF-dependent luciferase activity assay results (Figure 1B). We also observed a decrease in the protein levels of BNIP3L and BNIP3 in both cell lines and CAIX in U2OS cells when KDM2B was depleted under normal oxygen conditions (21% oxygen), indicating that loss of KDM2B reduces HIF activity in both normal oxygen conditions and in hypoxia, although these changes were not statistically significant (*P*>0.05) (Figure 2A, Supplementary Figure S1). In HeLa HRE-luciferase cells, KDM2B depletion in hypoxia induced an increase in BNIP3 and BNIP3L protein levels (Supplementary Figure S2), in agreement with the HIF-dependent luciferase activity assay results (Figure 1B); however, these changes were not statistically significant (*P*>0.05) (Supplementary Figure S2). qPCR analysis confirmed successful siRNA-mediated depletion of KDM2B in U2OS and MDA-MB-231 cells (Figure 2B). HIFs regulate transcription by binding to HREs at the promoters and enhancers of their target genes [3]. We next determined if KDM2B depletion affects HIF-1α binding to its target genes. To this end, U2OS cells were depleted of KDM2B and exposed to hypoxia prior to ChIP-qPCR analysis (Figure 2C). In the absence of KDM2B, HIF-1α occupancy at the HREs of the HIF target genes *BNIP3*, *CA9*, and *PHD3 (EGLN3)* was reduced compared to control siRNA-treated cells, returning to the levels of control siRNA-treated cells kept at normal oxygen (Figure 2C). Taken together, these results demonstrate that KDM2B can function as a positive regulator of HIF activity and recruitment of HIF-1α to its target gene HREs in certain cell types.

**Figure 2.**
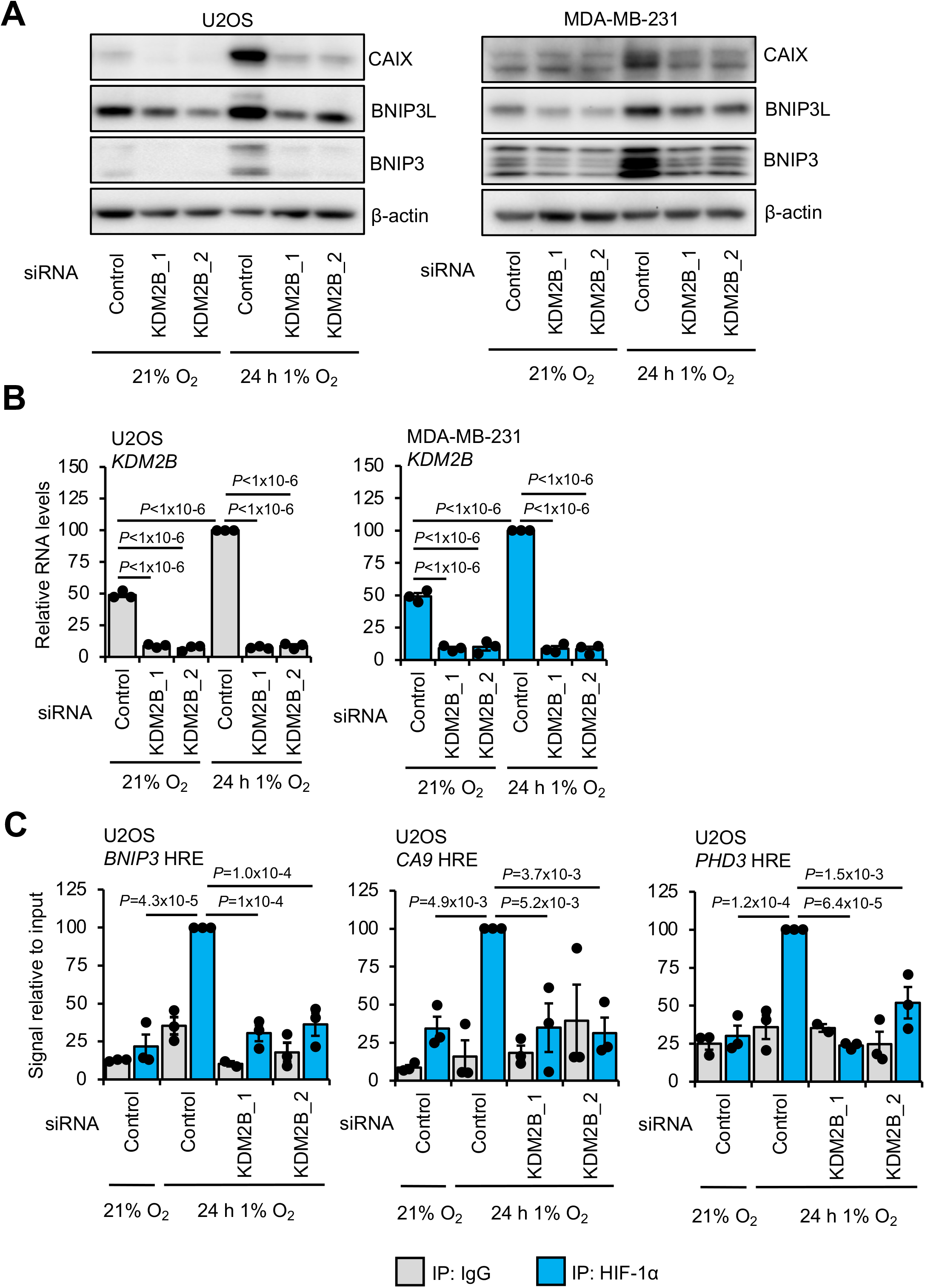
KDM2B regulates HIF activity. **A)** Western blot analysis for the indicated proteins in U2OS and MDA-MB-231 cells cultured at 21% oxygen, transfected with control, KDM2B_1, or KDM2B_2 siRNA for 48 hours, and exposed or not to 1% oxygen for 24 hours. A representative Western blot of three biological replicates is shown. **B)** qPCR analysis of *KDM2B* RNA levels in U2OS and MDA-MB-231 cells cultured at 21% oxygen, transfected with control, KDM2B_1, or KDM2B_2 siRNA for 48 hours, and exposed or not to 1% oxygen for 24 hours. *ACTB* was used as a normalising gene. Data represent mean (n=3 biological replicates) ± SEM. *P* values for group comparisons were calculated by one-way ANOVA with Tukey’s test. **C)** ChIP-qPCR analysis in U2OS cells cultured at 21% oxygen, transfected with control, KDM2B_1, or KDM2B_2 siRNA for 48 hours, and exposed or not to 1% oxygen for 24 hours. ChIP was performed with HIF-1α and non-specific antibody (IgG) immunoprecipitations (IPs), followed by DNA extraction and qPCR analysis at hypoxia response elements (HREs) located on the promoters of *BNIP3*, *CA9*, and *PHD3 (EGLN3)*. Data represent mean (n=3 biological replicates) ± SEM. *P* values for group comparisons were calculated by one-way ANOVA with Dunnett’s test.

### KDM2B is required for full HIF-1α expression

As we observed reduced HIF activity and recruitment to target genes upon KDM2B loss, we next assessed the consequences of KDM2B loss on HIF levels (Figure 3, Supplementary Figure S3). As expected, HIF-1α and HIF-2α protein levels were elevated in response to hypoxia exposure (24 hours 1% oxygen) in control siRNA-treated cells (Figure 3A, Supplementary Figure S3). KDM2B depletion decreased HIF-1α protein levels in both control oxygen (21% oxygen) and hypoxia-exposed U2OS and MDA-MB-231 cells (Figure 3A, Supplementary Figure S3). No consistent change in HIF-1β or HIF-2α protein levels was observed when KDM2B was depleted in U2OS or MDA-MB-231 cells (Figure 3A, Supplementary Figure S3), although HIF-2α protein levels did increase in hypoxia (*P*>0.05) with one of two KDM2B siRNAs. Our previous work has highlighted that HIFs can be regulated at the transcriptional level, resulting in changes in protein and activity [11, 14, 30]. As such, and since KDM2B is a chromatin-modifying protein, we next investigated if KDM2B could be controlling the mRNA levels of HIF-1α. From qPCR analysis, KDM2B depletion led to a reduction in *HIF1A* mRNA levels in U2OS and MDA-MB-231 cells, independent of hypoxia exposure (Figure 3B). No significant changes were observed in *HIF2A* (*EPAS1*) RNA upon KDM2B depletion in either cell line, although there was an upward trend in U2OS cells (Supplementary Figure S4). These data show that KDM2B is important for maintaining HIF-1α levels both in hypoxia and under control oxygen tensions. Furthermore, the reduction in HIF-dependent luciferase activity, HIF target gene expression, and HIF-1α occupancy at its target gene promoters when KDM2B is depleted is likely through reduced HIF-1α gene expression. We also assessed the effect of KDM2B depletion on HIF protein levels in HeLa cells and found that HIF-1α protein levels in hypoxia are elevated with one KDM2B siRNA and HIF-2α protein levels in hypoxia are elevated with both KDM2B siRNAs (Supplementary Figure S5). The increase in HIF-dependent reporter activity (Figure 1B) and increase in HIF target gene expression (Supplementary Figure S2), when KDM2B is depleted in HeLa cells, is likely explained by the increase in HIF-α protein levels. The contrasting results in HeLa cells compared to U2OS and MDA-MB-231 cells again highlight the context-dependent nature of HIF regulation by KDM2B.

**Figure 3.**
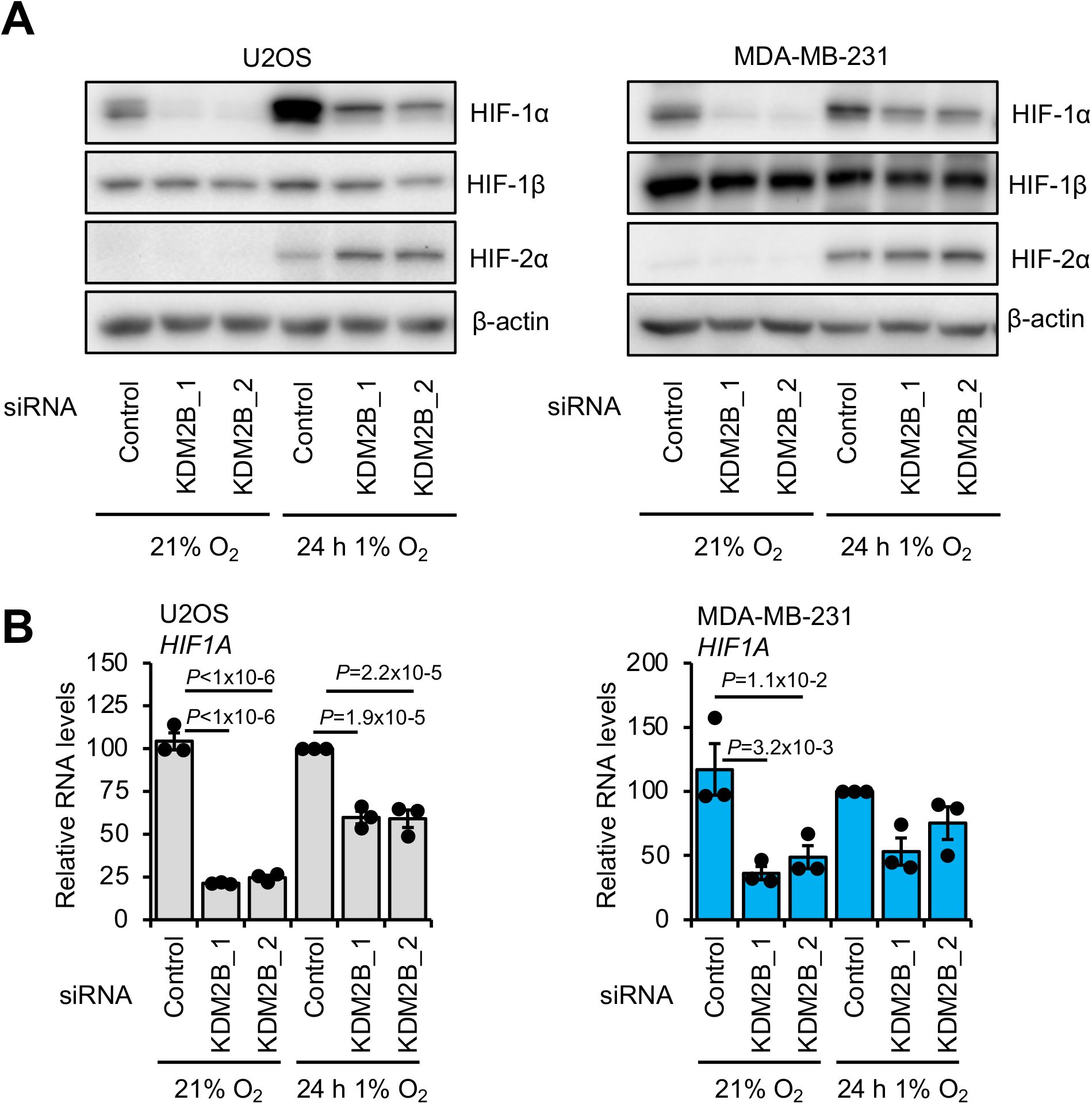
KDM2B regulates HIF-1α mRNA and protein expression. **A)** Western blot analysis for the indicated proteins in U2OS and MDA-MB-231 cells cultured at 21% oxygen, transfected with control, KDM2B_1, or KDM2B_2 siRNA for 48 hours, and exposed or not to 1% oxygen for 24 hours. A representative Western blot of three biological replicates is shown. **B)** qPCR analysis of *HIF1*A RNA levels in U2OS and MDA-MB-231cells cultured at 21% oxygen, transfected with control, KDM2B_1, or KDM2B_2 siRNA for 48 hours, and exposed or not to 1% oxygen for 24 hours. *ACTB* was used as a normalising gene. Data represent mean (n=3 biological replicates) ± SEM. *P* values for group comparisons were calculated by one-way ANOVA with Tukey’s test.

### KDM2B requires its JmjC and CxxC domains for positive regulation of HIF

KDM2B is a complex protein, with several functional domains [26, 28]. This includes a JmjC domain for catalytic demethylase activity, a CxxC domain important for DNA binding, a Plant Homo Domain (PHD), which is a chromatin reader, an F-box domain, which is a component of the SCF ubiquitin ligase complex, and leucine rich repeats (LRRs) (FBOX associated domain). Thus, to gain mechanistic insight, we elucidated the contribution of these domains to the control of HIF activity and expression by KDM2B (Figure 4). To this end, we generated and utilised loss-of-function mutant plasmids for KDM2B’s JmjC, CxxC, PHD, and F-Box domains (Figure 4A). We assessed the importance of each of these domains for the control of HIF-dependent reporter activity by KDM2B by comparing KDM2B wildtype and mutant overexpression in U2OS HRE-luciferase cells under control oxygen tensions (21% oxygen) (Figure 3B). Here, we could observe, as also seen in Figure 1D, that transient overexpression of KDM2B wildtype increased HIF-dependent luciferase activity, further supporting KDM2B’s function as a positive regulator of HIF activity (Figure 3B). With loss of the JmjC demethylase and CxxC DNA binding domain functions, KDM2B overexpression no longer resulted in increased HIF-dependent luciferase activity (Figure 4B). On the other hand, loss of PHD or F-box domain function had no effect on HIF-dependent luciferase activity when compared to wildtype KDM2B. Due to difficulties in overexpressing KDM2B in MDA-MB-231 cells, we repeated these experiments in HEK293 HRE-luciferase cells, and similar results were found in this cell line (Figure 4B). We also performed Western blot analysis in U2OS and HEK293 HRE-luciferase cells overexpressing KDM2B wildtype and mutant plasmids (Figure 4C, Supplementary Figure S6-S7). KDM2B was robustly overexpressed to similar levels in cells transfected with KDM2B wildtype and KDM2B domain mutant plasmids, demonstrating successful transfection and showing that these domain mutants do not affect KDM2B protein expression compared to the wildtype (Figure 4C, Supplementary Figure S6-S7). Protein levels of HIF-1α and the HIF target genes, CAIX (gene name *CA9*) and BNIP3, increased in KDM2B wildtype, KDM2B PHD, and KDM2B F-Box loss-of-function overexpressing U2OS HRE-luciferase cells, but not in KDM2B JmjC and CxxC loss-of-function overexpressing U2OS HRE-luciferase cells (Figure 4C, Supplementary Figure S6). Similar results were observed in HEK293 HRE-luciferase cells (Figure 4C, Supplementary Figure S7); however, in this cell line, the changes in HIF-1α protein levels were not statistically significant (*P*>0.05) (Figure 4C, Supplementary Figure S7), and we could not detect CAIX by Western blotting under control oxygen conditions in this cell line. Lastly, to complement the protein analysis, we also measured *HIF1A* RNA levels in response to overexpression of KDM2B wild type and domain loss-of-function plasmids in U2OS HRE-luciferase cells (Figure 4D). Similar to HIF-1α protein levels, *HIF1A* was increased in cells overexpressing KDM2B wild type and PHD and F-Box loss-of-function mutants, and this response was lost in cells overexpressing KDM2B JmjC and CxxC loss-of-function mutants (Figure 4D). These results provide further evidence for KDM2B’s function as a positive regulator of HIF activity and indicate that mechanistically, regulation of HIF activity and HIF-1α expression is dependent on KDM2B’s demethylase activity and DNA binding ability.

**Figure 4.**
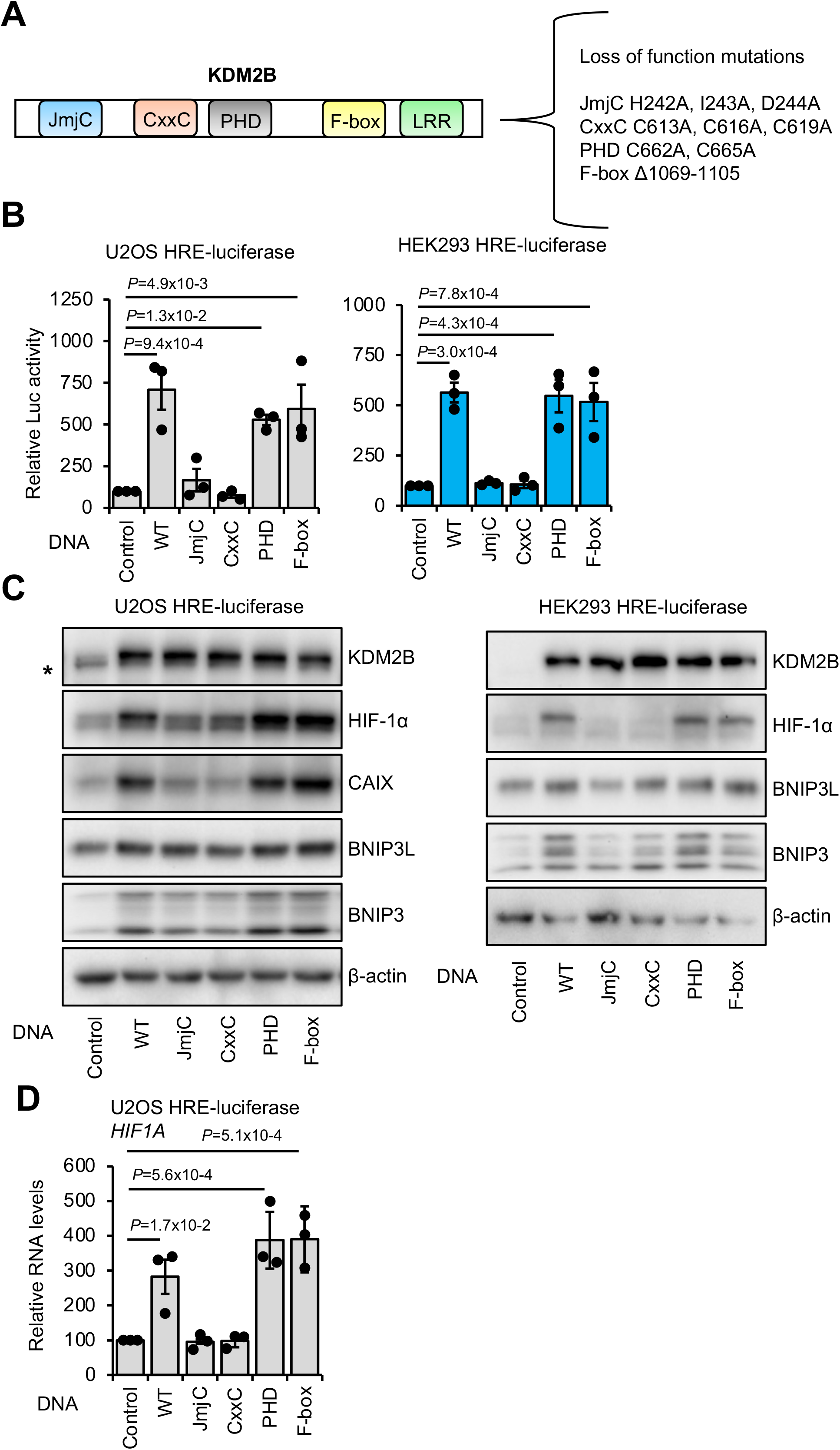
KDM2B controls HIF activity through its JmjC and CxxC domains. **A)** Schematic of KDM2B protein domains and loss-of-functions mutations used in this study. **B)** Luciferase assay analysis in U2OS and HEK293 HRE-luciferase cells cultured at 21% oxygen and transfected with control, KDM2B WT, KDM2B JmjC, KDM2B CxxC, KDM2B PHD, or KDM2B F-box plasmids for 48 hours. Data represent mean (n=3 biological replicates) ± SEM. *P* values for group comparisons were calculated by one-way ANOVA with Dunnett’s test. **C)** Western blot analysis for the indicated proteins in U2OS and HEK293 HRE-luciferase cells cultured at 21% oxygen and transfected with control, KDM2B WT, KDM2B JmjC, KDM2B CxxC, KDM2B PHD, or KDM2B F-box plasmids for 48 hours. A representative Western blot of three biological replicates is shown. * Lower band is non-specific. **D)** qPCR analysis of HIF1A RNA levels in U2OS HRE-luciferase cells cultured at 21% oxygen and transfected with control, KDM2B WT, KDM2B JmjC, KDM2B CxxC, KDM2B PHD, or KDM2B F-box plasmids for 48 hours. *ACTB* was used as a normalising gene. Data represent mean (n=3 biological replicates) ± SEM. *P* values for group comparisons were calculated by one-way ANOVA with Dunnett’s test.

### KDM2B mediates RNA Pol II recruitment at the *HIF1A* promoter

Although KDM2B predominantly functions as a transcriptional repressor, via Polycomb repressive dependent and independent mechanisms [31], it can also activate transcription [32–34], including, in the case of *IL6*, through RNA Pol II recruitment [33]. To begin to elucidate the molecular mechanisms by which KDM2B controls HIF-1α RNA expression, we performed ChIP-qPCR analysis to measure RNA Pol II recruitment to the *HIF1A* promoter (Figure 5A). Loss of KDM2B through siRNA depletion in U2OS cells reduced RNA Pol II binding to the *HIF1A* promoter, both in control oxygen levels (21% oxygen) and in response to hypoxia (24 hours 1% oxygen) (Figure 5B). Conversely, overexpression of wild type KDM2B in U2OS cells increased *HIF1A* promoter RNA Pol II occupancy (Figure 5C). Upregulated RNA Pol II recruitment remained in cells overexpressing KDM2B with PHD or F-Box loss-of-function mutants, but overexpression of KDM2B with JmjC or CxxC domain mutants had no effect on RNA Pol II recruitment (Figure 5C). Western blot analysis confirmed KDM2B wild type and mutant overexpression in U2OS cells (Figure 5 D-E). These data indicate that KDM2B’s regulation of HIF-1α expression occurs, at least in part, through controlling RNA Pol II occupancy at the *HIF1A* promoter, via its JmjC and CxxC domains.

**Figure 5.**
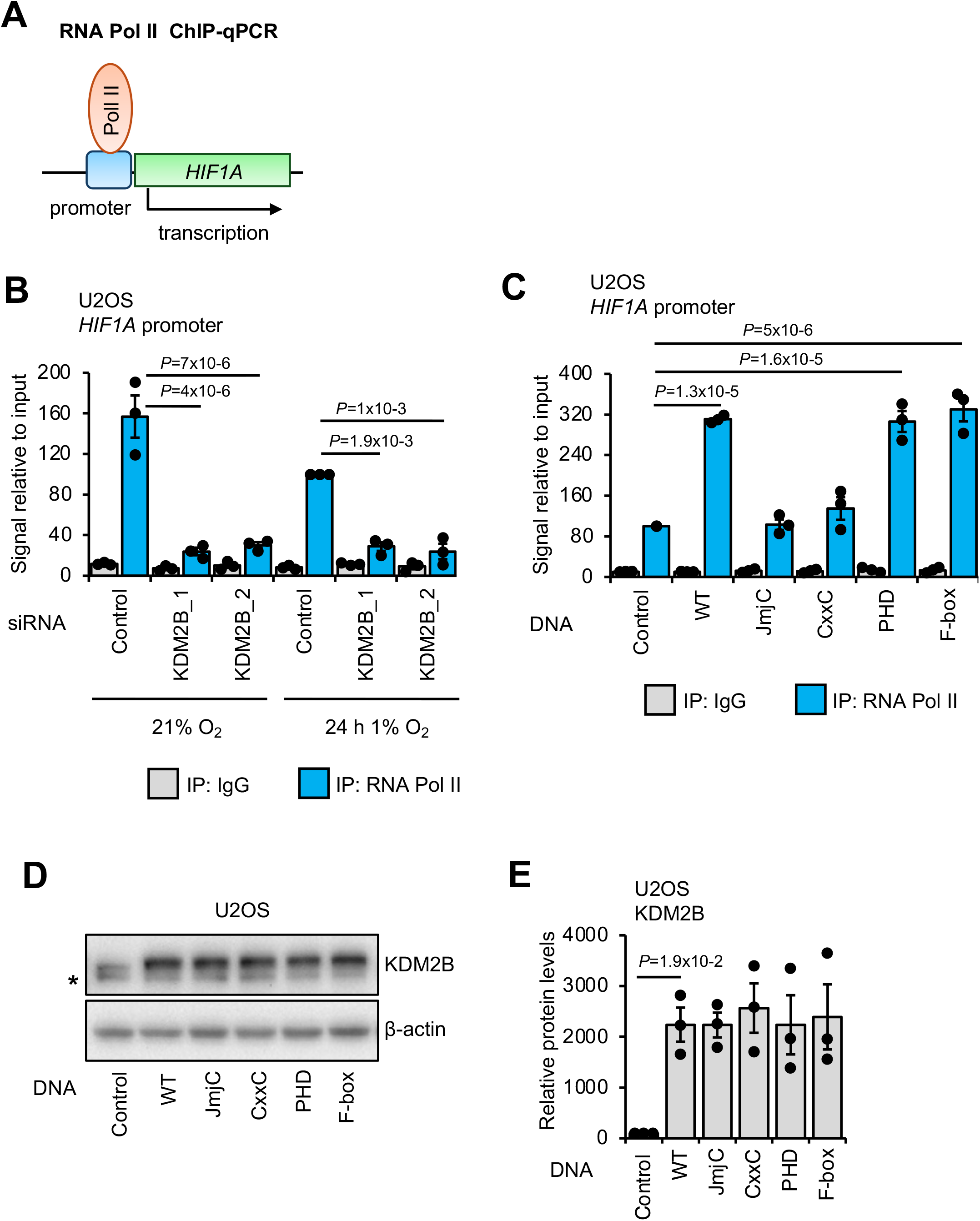
Recruitment of RNA Pol II to the *HIF1A* promoter is modulated by KDM2B. **A)** Schematic of ChIP-qPCR analysis to measure binding of RNAP2 to the *HIF1A* promoter. **B)** ChIP-qPCR analysis U2OS cells cultured at 21% oxygen, transfected with control, KDM2B_1, or KDM2B_2 siRNA for 48 hours, and exposed or not to 1% oxygen for 24 hours. ChIP was performed with RNA Pol II and non-specific antibody (IgG) immunoprecipitations (IPs), followed by DNA extraction and qPCR analysis at the *HIF1A* promoter. Data represent mean (n=3 biological replicates) ± SEM. *P* values for group comparisons were calculated by one-way ANOVA with Tukey’s test. **C)** ChIP-qPCR analysis in U2OS cells cultured at 21% oxygen and transfected with control, KDM2B WT, KDM2B JmjC, KDM2B CxxC, KDM2B PHD, or KDM2B F-box plasmids for 48 hours. ChIP was performed with RNA Pol II and non-specific antibody (IgG) immunoprecipitations (IPs), followed by DNA extraction and qPCR analysis at the *HIF1A* promoter. Data represent mean (n=3 biological replicates) ± SEM. *P* values for group comparisons were calculated by one-way ANOVA with Dunnett’s test. **D)** Western blot and **E)** Western blot quantification analysis for KDM2B protein levels in U2OS cells cultured at 21% oxygen and transfected with control, KDM2B WT, KDM2B JmjC, KDM2B CxxC, KDM2B PHD, or KDM2B F-box plasmids for 48 hours. β-actin was used as a normalising protein. A representative Western blot of three biological replicates is shown. Data represent mean (n=3 biological replicates) ± SEM. *P* values for group comparisons were calculated by one-way ANOVA with Dunnett’s test. * Lower band is non-specific.

We have previously shown that NF-κB controls *HIF1A* mRNA levels and HIF activity [11]. To initially investigate a potential link between NF-κB and the regulation of HIF by KDM2B, we depleted KDM2B in U2OS κB-luciferase cells and activated the NF-κB response with TNF-α stimulation (10 ng/µL TNF-α for 8 hours) (Supplementary Figure S8A). siRNA-mediated depletion of KDM2B resulted in lower NF-κB-dependent luciferase activity in U2OS κB-luciferase cells following TNF-α stimulation (Supplementary Figure S5A). Conversely, overexpression of KDM2B resulted in higher NF-κB-dependent luciferase activity following TNF-α stimulation (Supplementary Figure S5B). In a similar manner to HIF regulation, KDM2B regulation of NF-κB required functional JmjC and CxxC domains (Supplementary Figure S5B). This preliminary finding demonstrates that KDM2B can control NF-κB activity through its demethylase activity and DNA binding ability and provides a potential link to KDM2B-mediated control of HIF-1α expression.

### KDM2B depletion results in reduced proliferation in control and hypoxic conditions

A range of cellular responses are triggered in hypoxia, one of which is cell proliferation. To determine the biological significance of KDM2B depletion in cells in hypoxia, we measured cell proliferation over several days (Figure 6). As we have previously shown [30, 35], 1% oxygen treatment reduced proliferation in U2OS but also in MDA-MB-231 cells (Figure 6A-B). Under normal oxygen levels (21% oxygen), cells depleted of KDM2B proliferated significantly less compared to control siRNA-treated cells (Figure 6A-B). Interestingly, even in hypoxia, depletion of KDM2B resulted in a significantly reduced cell number compared to control siRNA-treated cells, suggestive of cell death (Figure 6A-B), particularly in U2OS cells, where numbers went below the initial number seeded. These findings show that KDM2B, further to its control of HIF-1α expression and HIF activity, is required for cellular responses in hypoxia.

**Figure 6.**
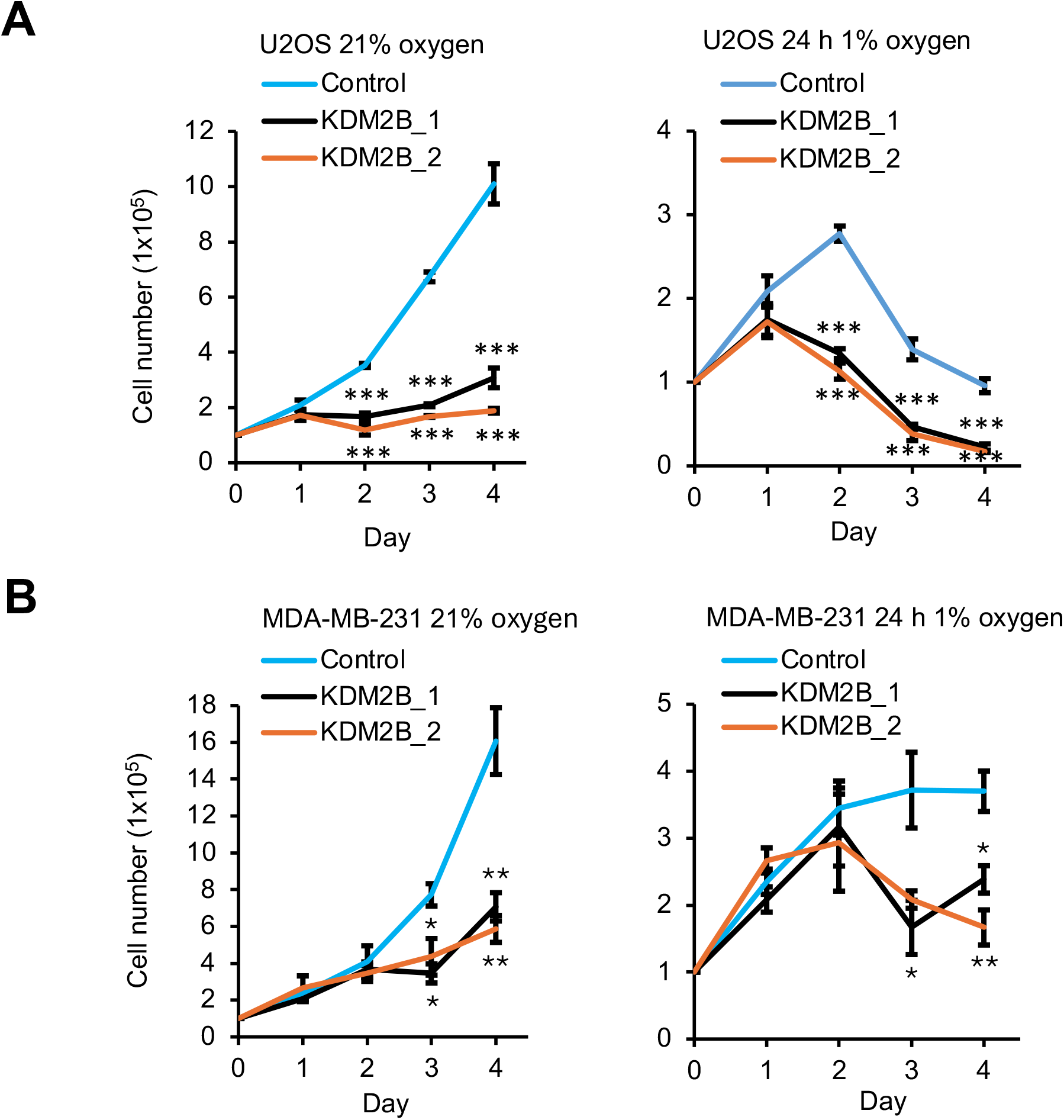
Cell proliferation is impaired in response to KDM2B depletion in normal oxygen conditions and hypoxia. Cell proliferation assay in **A)** U2OS and **B)** MDA-MB-231 cells cultured at 21% oxygen, transfected with control, KDM2B_1, or KDM2B_2 siRNA for 96 hours, and exposed or not to 1% oxygen for 72 hours. Data represent mean (n=3 biological replicates) ± SEM. *P* values for group comparisons were calculated by one-way ANOVA with Dunnett’s test.

Given the importance of KDM2B and HIF in cancer, and our evidence for the role of KDM2B as a positive regulator of HIF-1α expression and activity, we assessed the correlation of *HIF1A* and *KDM2B* mRNA levels in patient samples across different cancers from The Cancer Genome Atlas (TCGA) (Supplementary Figure S9). This analysis revealed that several human cancer types have a moderate positive (0.3-0.5) correlation that is highly statistically significant between levels of *KDM2B* and *HIF1A* mRNA (Supplementary Figure S9A), further supporting our mechanistic findings, whereby in most of the cell lines tested, KDM2B acts as a positive regulator of HIF-1α expression. We also show the correlation plots for BRCA (Breast Invasive Carcinoma) and CESC (Cervical Squamous Cell Carcinoma and Endocervical Adenocarcinoma), the most relevant to the cancer cell lines used in this study (Supplementary Figure S9B). Lastly, we also assessed the correlation between *KDM2B* RNA levels and the RNA levels of hypoxia gene signatures (Buffa [36], Ragnum [37], and Winter [38]) in breast and cervical cancer samples from The Cancer Genome Atlas (TCGA) (Supplementary Figure S10). Positive correlations between *KDM2B* and all three hypoxia gene signatures were present in the Breast Invasive Carcinoma data (Supplementary Figure S10A), complementing our findings in MB-MDA-231 cells.

## Discussion

Here, we show that in several cell types, KDM2B is required for full HIF-1α levels and activity. We find that depletion of KDM2B results in lower levels of HIF-1α mRNA and hence protein levels. This is accompanied by reduced occupancy of HIF-1α at HREs of its target genes and decreased expression of HIF target genes. Control of HIF-1α and HIF target gene expression, although more pronounced under hypoxia, was also present in normal oxygen conditions, indicating a mechanism independent of oxygen sensing. Mechanistically, we determined that KDM2B’s regulation of HIF-1α expression and HIF activity is dependent on its DNA binding capability (CxxC domain) and demethylase activity (JmjC domain). We also demonstrate that KDM2B mediates RNA Pol II to the *HIF1A* promoter. Finally, we show that KDM2B is required for maintaining cell proliferation under normal oxygen tensions and in hypoxia. Interestingly, in HeLa cells, we find that loss of KDM2B increases HIF activity and HIF-2α expression in hypoxia. This demonstrates that KDM2B-mediated HIF regulation is context dependent; however, presently we do not have a mechanistic insight into this cell-context dependence.

Although HIF-1α regulation is canonically regulated post-translationally via the activity of PHDs [39], we and others have demonstrated important mechanisms controlling HIF-1α gene expression [11, 12, 30], mRNA stability [40], and translation [41, 42]. All these mechanisms precede the ability of PHDs to target HIF-1α or HIF-1α stabilisation, and as such are as important in controlling HIF levels and activity. These findings add a new layer to the regulation of HIF-1α gene expression.

KDM2B has several functional domains. KDM2B can control transcription through its CxxC domain, independent of its demethylase activity, through recruitment of PRC1 to CpG islands [43–46], and through constraining RNA Pol II binding at CpG island promoters [31]. KDM2B can also act directly as an epigenetic silencer through its demethylase activity [28] and can target c-Fos for polyubiquitylation and proteasomal degradation through its F-Box domain [47]. In this study, we find that KDM2B controls HIF-1α expression and HIF activity through its CxxC domain and JmjC domain, and independent of its PHD domain and F-Box domain.

Here we find that KDM2B positively regulates *HIF1A* expression and recruitment of RNA Pol II to the *HIF1A* promoter. KDM2B has previously been shown to function as a transcriptional activator [32–34] and directly recruit BRG1 and RNA Pol II to the *IL6* gene promoter [33]. However, it is not clear from our work if KDM2B is acting directly on the *HIF1A* gene as a transcriptional activator or indirectly through other chromatin-binding proteins or transcription factors. Our own previous work has highlighted the important roles of NF-κB and SWI/SNF in controlling *HIF1A* mRNA levels in cells from several organisms [11, 12, 30]. Our preliminary work here demonstrates that KDM2B can also positively regulate NF-κB activity in TNF-α stimulated cells, and this regulation is dependent on the demethylase (JmjC) and DNA binding (CxxC) domains of KDM2B. KDM2B has been shown to regulate NF-κB in certain systems, such as HK2 cells [48]. However, a previous study in myocardial ischemia-reperfusion injury revealed that KDM2B depletion resulted in increased levels of NF-κB p65 and some of its genes [49]. Two separate studies demonstrated that KDM2B can bind BRG1 (core component of SWI/SNF) at some of the genes studied [33, 50]. These findings support a hypothesis that KDM2B is required for NF-κB-dependent control of HIF-1α gene expression, a process we have shown to be BRG1-dependent [11, 30]. Therefore, our data demonstrating that KDM2B is required for *HIF1A* mRNA levels agree with these previous links to NF-κB and SWI/SNF. However, additional investigation is needed to fully delineate this possibility. Interestingly, KDM2B KO embryos were shown to lack vasculature [51], an aspect also controlled by HIF-1α during development [52–54], further supporting our observations from this study.

KDM2B has recognised roles in promoting proliferation across a variety of cancer types. However, the mechanisms vary significantly, regulating Wnt [55], Myc [34], and PI3K [56, 57] pathways. Our results support all previous studies, demonstrating that KDM2B depletion leads to reduced proliferation. In addition, we have shown that KDM2B is required to maintain cell numbers in hypoxia. It is probable that, given the reduction in cell survival pathways such as PI3K, Myc, and HIF, when KDM2B is depleted, cells under such stress as hypoxia will not be able to survive. Furthermore, KDM2B has been shown to control autophagy [56, 58], a process that hypoxic cells induce via BNIP3 and BNIP3L [35, 59], used for survival. Our results show that BNIP3 levels are reduced when KDM2B is depleted, suggesting autophagy induction in hypoxia is impaired. This would result in reduced survival, much like the reduction in cell numbers we observed in cells depleted of KDM2B and exposed to hypoxia. In lung [56] and gastric cancer [58], depletion of KDM2B induced autophagy, a process dependent on the inhibition of mTOR. Interestingly, Xie et al [56] also reported a reduction of several HIF-1α genes involved in glycolysis in lung cancer cells when KDM2B is depleted. These included Glut1 and LDHA; however, this study did not investigate HIF-1α levels, focusing on mTOR, a known regulator of HIF-1α [60, 61] and NF-κB [62]. We also find that *KDM2B* and *HIF1A* RNA expression positively correlate across a variety of cancer types from The Cancer Genome Atlas (TCGA), and that *KDM2B* and hypoxia gene signature RNA expression positively correlate in breast invasive carcinoma samples from The Cancer Genome Atlas (TCGA), further supporting our mechanistic findings that KDM2B promotes HIF-1α expression and HIF activity, including in breast cancer cells.

We have previously shown that the KDM2 subfamily (KDM2A and KDM2B) of JmjC domain containing proteins are HIF-1 target genes [25]. This upregulation of KDM2B in hypoxia may act as a positive feedback mechanism to maintain HIF activity in certain contexts. Future work should assess the potential role of KDM2A in controlling HIF activity and the hypoxia response. KDM2B demethylates H3K36me1/2 and H3K4me3 through its JmjC domain, and we find that KDM2B requires its histone demethylase activity for modulation of HIF. However, we did not examine the effects of KDM2B on the histone methylation status genome-wide, or at the HIF-1α gene. JmjC domain containing proteins require oxygen for their enzymatic activity, and oxygen-sensitive impairment of JmjC protein histone demethylase activity in hypoxia has been reported for KDM5A [18], KDM6A [19], and KDM4A [63]. Here we report that loss of KDM2B under control oxygen conditions (21% oxygen) and hypoxia (24 hours of 1% oxygen), reduces HIF activity and HIF-1α expression. This suggests that the histone demethylase activity of KDM2B is retained at 1% oxygen in cells, as is seen with KDM3A and KDM4C, which are both HIF target genes that remain active at low oxygen concentrations and function in hypoxia as HIF coactivators through their histone demethylase activity [22–24]. Further investigation is required to determine the role of KDM2B-mediated histone demethylation in the context of HIF regulation and the hypoxia response.

Taken together, our results indicate that KDM2B is a positive regulator of HIF-1α expression and HIF activity in several cell types, and this is important for maintaining cell numbers in hypoxia. Further mechanistic investigation on how KDM2B controls the *HIF1A* gene will be necessary to fully understand this phenomenon. Furthermore, using KDM2B inhibitors might result in reduced HIF signalling, as an advantage for certain pathologies such as cancer, but be detrimental for others such as neurodegeneration, where HIF-mediated survival pathways might be needed.

## Methods

### Reagents and tools table

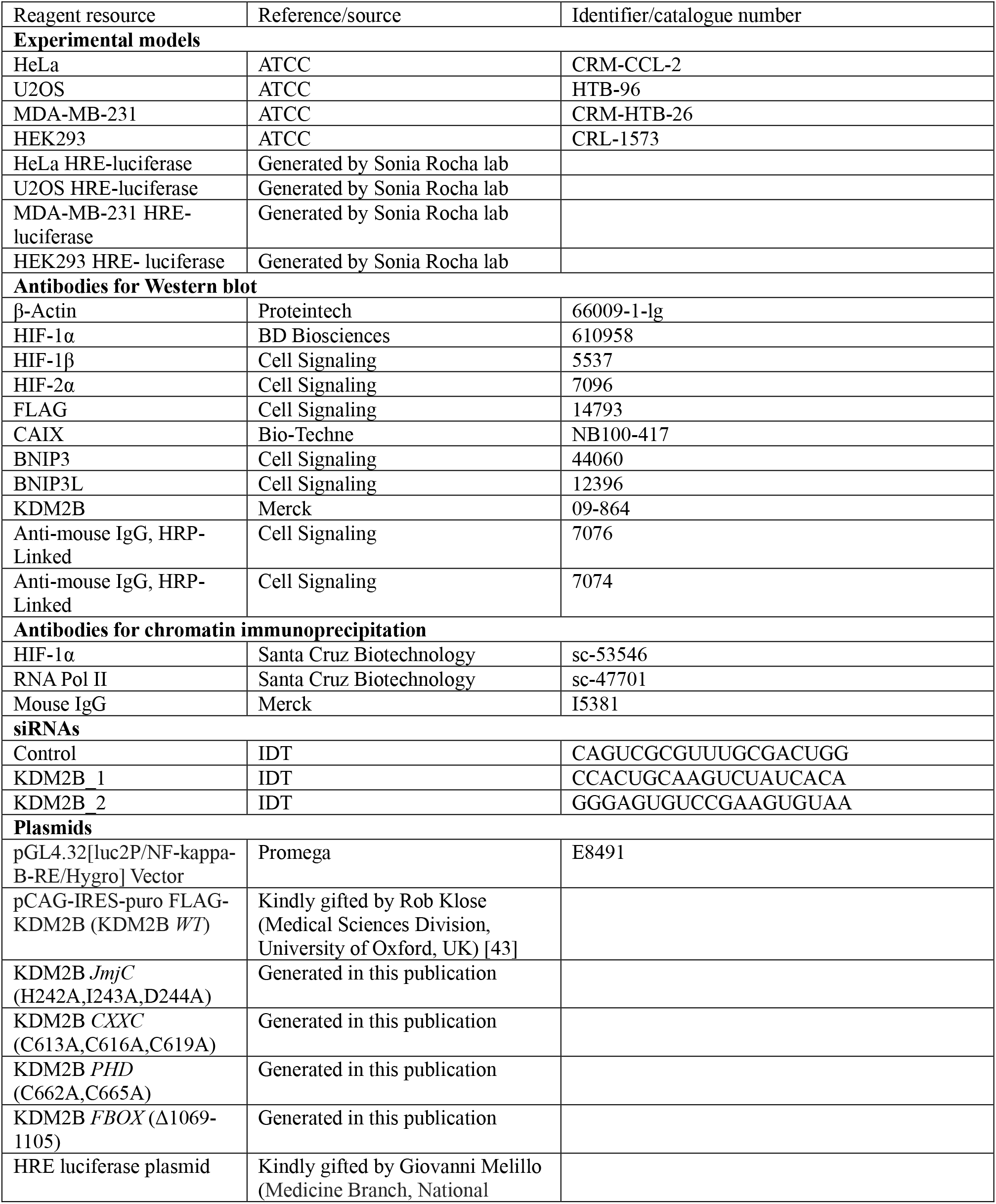
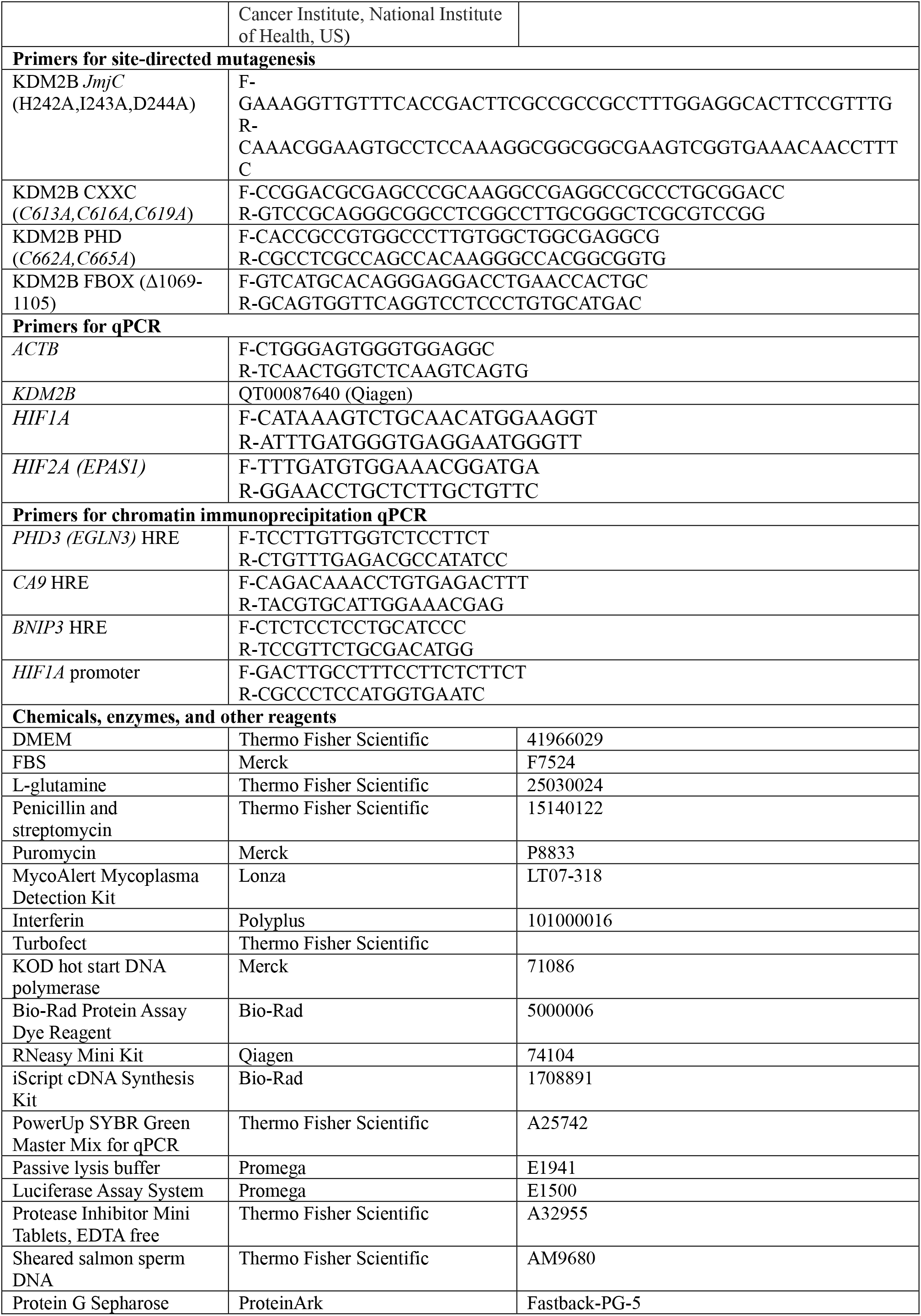
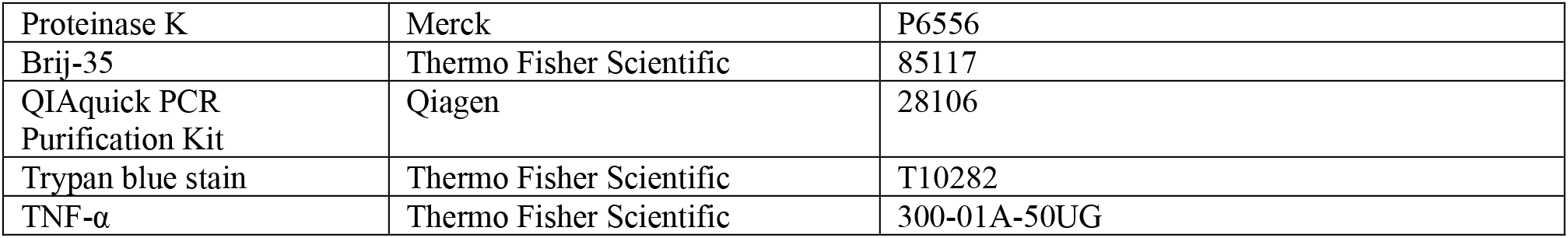

### Cell lines

HeLa, U2OS, and MDA-MB-231 cell lines were maintained in DMEM supplemented with 10% v/v FBS, 2 mM L-glutamine, 100 units/mL penicillin, and 100 µg/mL streptomycin. U2OS-HRE (hypoxia response element), MDA-MB-231-HRE, HeLa-HRE, and HEK293-HRE luciferase lines were generated in the lab by co-transfection with a HRE luciferase plasmid containing 3 copies of 21 bps of the iNOS 21bp HRE (5’-AGTGACTACGTGCTGCCTAGG-‘3) subcloned into KpnI and MluI restriction sites of the pGL3 promoter vector (Promega), which was kindly gifted by G. Melillo (National Cancer Institute, National Institutes of Health, Bethesda, MD), together with a puromycin-resistance gene containing vector. U2OS-κB luciferase cells were generated in the lab by transfection with the pGL4.32(luc2P/NF-κB-RE/Hygro) vector. Additionally, 0.5μg/mL puromycin was added to cell culture media for U2OS-HRE, MDA-MB-231-HRE, and HEK293-HRE luciferase cell lines, and 150 μg/mL hygromycin for U2OS-κB luciferase cells. Cell lines were cultured for no more than 30 passages and routinely tested for mycoplasma contamination using a MycoAlert Mycoplasma Detection Kit.

### Cell treatments

Hypoxia treatments were performed by incubating cells in an Invivo2 300 hypoxia workstation (Baker Ruskin) at 1% O_2_, 5% CO_2_, and 37 °C. To avoid reoxygenation, cells were lysed in the workstation. TNF-α (Peprotech) was dissolved in PBS and used at a final concentration of 10 ng/mL.

### siRNA transfections

HEK293 cells were transfected in 35mm plates with a final concentration of 27 nM of siRNA by mixing siRNA, 0.12 M CaCl2, HEPES buffered saline (0.156 M NaCl, 0.375 M Na2HPO4, and 10mM HEPES), and water at a final volume of 400 µL and adding to cells. All other cell lines were transfected in 35mm plates with a final concentration of 27 nM of siRNA using Interferin (Poluplus, VWR) transfection reagent. For reverse transfections, the transfection mixture was added to the media immediately before cell seeding.

### DNA transfections

HEK293 cells were transfected in 35mm plates with 1 µg of DNA for 48 hours by mixing DNA, 0.12 M CaCl2, HEPES buffered saline (0.156 M NaCl, 0.375 M Na2HPO4, and 10mM HEPES), and water at a final volume of 400µL and adding to cells. All other cell lines were transfected in 35 mm plates with 1 µg of DNA for 48 hours using Turbofect transfection reagent. pCAGIpuro_hKDM2B CDS (a gift from R. Klose, Medical Sciences Division, University of Oxford, Oxford, UK) was used to generate KDM2B JmjC (H242A,I243A,D244A), CXXC (C613A,C616A,C619A), PHD (C662A,C665A), and FBOX (Δ1069-1105) domain mutants. HA-HIF1alpha-pcDNA3 was a gift from William Kaelin (Addgene plasmid # 18949).

### Site-directed mutagenesis

Site-directed mutagenesis was performed using a SureCycler 8800 thermocycler (Agilent) with the KOD hot start DNA polymerase kit. DNA sequencing was performed to confirm successful site-directed mutagenesis.

### Western blots

Cells were lysed in RIPA buffer (50 mM Tris-HCl pH 8.0, 150 mM NaCl, 1% v/v NP40, 0.25% w/v Na-deoxycholate, 0.1% w/v SDS, 10 mM NaF, 2 mM Na3VO4, and 1 protease inhibitor tablet per 10 mL of lysis buffer), incubated on ice for 10 minutes, and centrifuged for 15 minutes at 13,000 rpm and 4 °C. The supernatants (RIPA-soluble protein samples) were collected, and protein concentrations were determined by Bradford assay using Bio-Rad Protein Assay Dye Reagent. 20 µg of protein was prepared in 2 × SDS loading buffer (100 mM Tris-HCl pH 6.8, 20% v/v glycerol, 4% w/v SDS, 200 mM DTT and Bromophenol Blue). SDS-PAGE and Western blots were carried out using standard protocols. Images were obtained by enhanced chemiluminescence using a ChemiDoc (Bio-Rad).

### RNA expression analysis by qPCR

RNA was extracted using the RNeasy Mini Kit, and 300 ng of RNA was converted to cDNA using an iScript cDNA Synthesis Kit. qPCR was performed on a QuantStudio 1 qPCR platform (Applied Biosystems) with PowerUp SYBR Green Master Mix. The quantity of RNA was determined using the ΔΔCT method [64] and normalised to the reference gene *ACTB*.

### Luciferase assay

Cells were lysed in Passive lysis buffer. Luciferase activity was measured using a Luciferase Assay System kit and normalised to protein concentration determined by Bradford assay using Bio-Rad Protein Assay Dye Reagent.

### Chromatin immunoprecipitation (ChIP)-qPCR

Cells on 100 mm cell culture plates were crosslinked by adding formaldehyde at a final concentration of 1% v/v for 10 minutes. Crosslinking was performed inside the hypoxia chamber for hypoxia-exposed samples, or inside the cell culture incubator for control oxygen samples. Cells were quenched by adding glycine at a final concentration of 0.125 M for 10 minutes at room temperature. Cells were washed two times with cold PBS and lysed with 450 μL ChIP lysis buffer (1% w/v SDS, 10 mM EDTA pH 8, 50 mM Tris-HCl pH 8.1, and 1 protease inhibitor tablet) and incubated on ice for 10 minutes. Cell lysates were sonicated at 4 °C, 25 times for 15 seconds with 30 seconds gap between each sonication cycle at 50% amplitude using a Sonics Vibra Cell #VCX130 sonicator. Cell lysates were centrifuged for 15 minutes at 13,000 rpm and 4 °C. The supernatants (chromatin samples) were collected; 10 μL chromatin samples were collected as input samples (10% of immunoprecipitation input) and made up to 100 μL with water; 100 μL chromatin samples were diluted 10-fold with dilution buffer (1% v/v Triton X-100, 2 mM EDTA pH 8, 150 mM NaCl, and 20 mM Tris-HCl pH 8.1) and pre-cleared with 2 μg sheared salmon sperm DNA and 20 μL Protein G Sepharose (50% slurry with PBS) on a rotating wheel for 1 hour at 40 rpm and 4 °C. The diluted and pre-cleared chromatin samples were centrifuged for 1 minute at 1000 rpm and 4 °C. The supernatants were collected. Immunoprecipitations were performed by incubating the diluted and pre-cleared chromatin samples with protein target or control antibodies and 0.1% v/v Brij-35 rotating wheel overnight at 40 rpm and 4 °C. To capture immune complexes, 30 μL Protein G Sepharose (50% slurry with PBS) slurry and 2 μg sheared salmon sperm DNA were added to each immunoprecipitation sample, followed by incubation on a rotating wheel for 2 hours at 40 rpm and 4 °C, samples were centrifuged for 1 minute at 1000 rpm and 4 °C, and supernatants were discarded. The immune complexes (pellets) were resuspended in 1 mL of wash buffer 1 (0.1% w/v SDS, 1% v/v Triton X-100, 2 mM EDTA pH 8, 150 mM NaCl, and 20 mM Tris-HCl pH 8.1), incubated on a rotating wheel for 5 minutes at 40 rpm and 4 °C, centrifuged for 1 minute at 1000 rpm and 4 °C, and supernatants were discarded. The immune complexes (pellets) were resuspended in 1 mL of wash buffer 2 (0.1% w/v SDS, 1% v/v Triton X-100, 2 mM EDTA, 500 mM NaCl, and 20 mM Tris-HCl pH 8.1), incubated on a rotating wheel for 5 minutes at 40 rpm and 4 °C, centrifuged for 1 minute at 1000 rpm and 4 °C, and supernatants were discarded. The immune complexes (pellets) were resuspended in 1 mL of TE buffer (1 mM EDTA and 10 mM Tris-HCl pH 8.1), incubated on a rotating wheel for 5 minutes at 40 rpm, centrifuged for 1 minute at 1000 rpm, and supernatants were discarded. To elute the immune complexes, pellets were resuspended in 100 μL elution buffer (1% w/v SDS and 0.1 M NaHCO4), incubated on a rotating wheel for 1 hour at 40 rpm, centrifuged for 1 minute at 1000 rpm, and pellets were discarded. To reverse crosslinks, NaCl was added at a final concentration of 0.2 M to the input chromatin samples and eluted immunoprecipitated samples, which were incubated on a thermomixer for 4 hours at 500 rpm and 65 °C. To digest proteins, Proteinase K (20 μg), EDTA pH 8 (final concentration 10 mM), and Tris-HCl pH 6.5 (final concentration 40 mM) were added to each sample, and samples were incubated on a thermomixer for 1 hour at 500 rpm and 45 °C. DNA was purified using a QIAquick PCR Purification Kit. ChIP DNA was analysed using qPCR, performed on a QuantStudio 1 qPCR platform (Applied Biosystems) with PowerUp SYBR Green Master Mix.

### Cell proliferation assays

Cells were reverse transfected (see above section), and live cells were counted 24, 48, 72, and 96 hours post-transfection using a Countess 3 automated cell counter (Invitrogen) and trypan blue. Hypoxia exposure was started 24 hours after reverse transfection.

### *KDM2B* and *HIF1A* correlation analysis

Correlation analysis between *KDM2B* and *HIF1A* RNA expression in patient samples across different cancer types from The Cancer Genome Atlas (TCGA) was performed using GEPIA2 [65].

### *KDM2B* and hypoxia signature correlation analysis

Correlation analysis between *KDM2B* and hypoxia signature (Buffa [36], Ragnum [37], and Winter [38]) RNA expression in patient samples across different cancer types from The Cancer Genome Atlas (TCGA) PanCancer Atlas was performed using the clinical attribute feature of cBioPortal [66].

### Statistical Analysis

For qPCR, luciferase assays, and cell proliferation assays, a minimum of three biological replicates were analysed, and *P* values were calculated by one-way ANOVA with Tukey’s test for multiple pairwise group comparisons or one-way ANOVA with Dunnetts’s test for multiple pairwise group comparisons to one control group using GraphPad v10.4.0. For *KDM2B* and *HIF1A* RNA expression correlation analysis, *P* values and *R* values were calculated by Pearson’s correlation coefficient using GEPIA2 [65]. For *KDM2B* and hypoxia signature RNA expression correlation analysis, *P* values and *R* values were calculated by Pearson’s correlation coefficient using cBioPortal [66].

## Data availability

Source data is available at Figshare: https://doi.org/10.6084/m9.figshare.33154961 [67].

## Acknowledgements

We would like to thank Dr. James W Wilson, Mr Nathan Williams and Mr Orran Simpson for technical discussions during this work. We would like to thank Rob Klose (Oxford University) for the KDM2B expression plasmid.

## Author contributions

CRediT Author Contribution = Sonia Rocha: Conceptualization, Supervision, Funding acquisition, Writing - Review & Editing. Michael Batie: Conceptualization, Formal analysis, Validation, Investigation, Visualization, Methodology, Writing - Original Draft, Writing - Review & Editing. Dilem Shakir: Investigation, Visualization, Chun-Sui Kwok: Investigation, Gemma Bell: Investigation, Jiahua Kou: Investigation, Ali Bakhsh: Investigation.

