## Supplementary data for "KDM2B controls HIF levels and activity through its JmjC and CxxC domains"

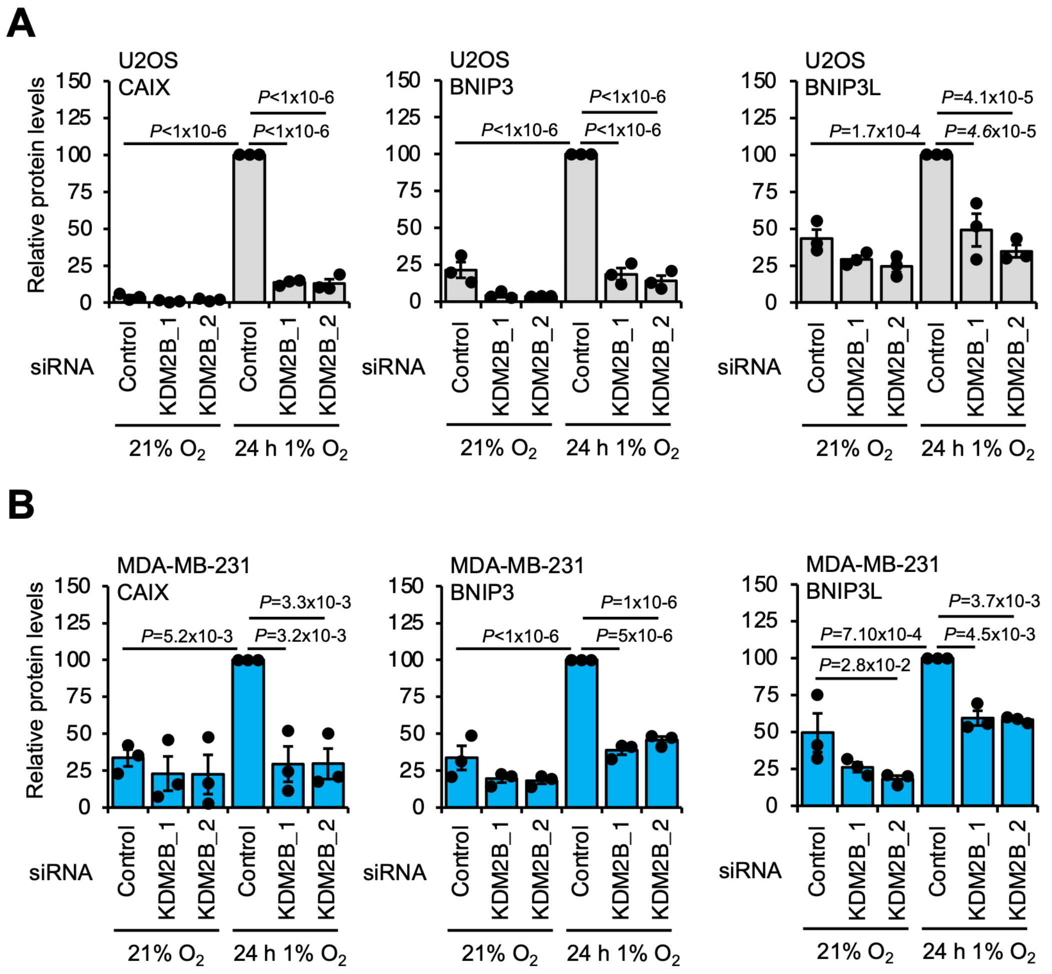


**Supplementary Figure S1. Regulation of HIF activity by KDM2B in U2OS and MDA-MB-231 cells, Western blot quantification**. Western blot quantification analysis for the indicated proteins in **A)** U2OS and **B)** MDA-MB-231 cells cultured at 21% oxygen, transfected with control, KDM2B_1, or KDM2B_2 siRNA for 48 hours, and exposed or not to 1% oxygen for 24 hours. β-actin was used as a normalising protein. Data represent mean (n=3 biological replicates) ± SEM. *P* values for group comparisons were calculated by one-way ANOVA with Tukey's test.

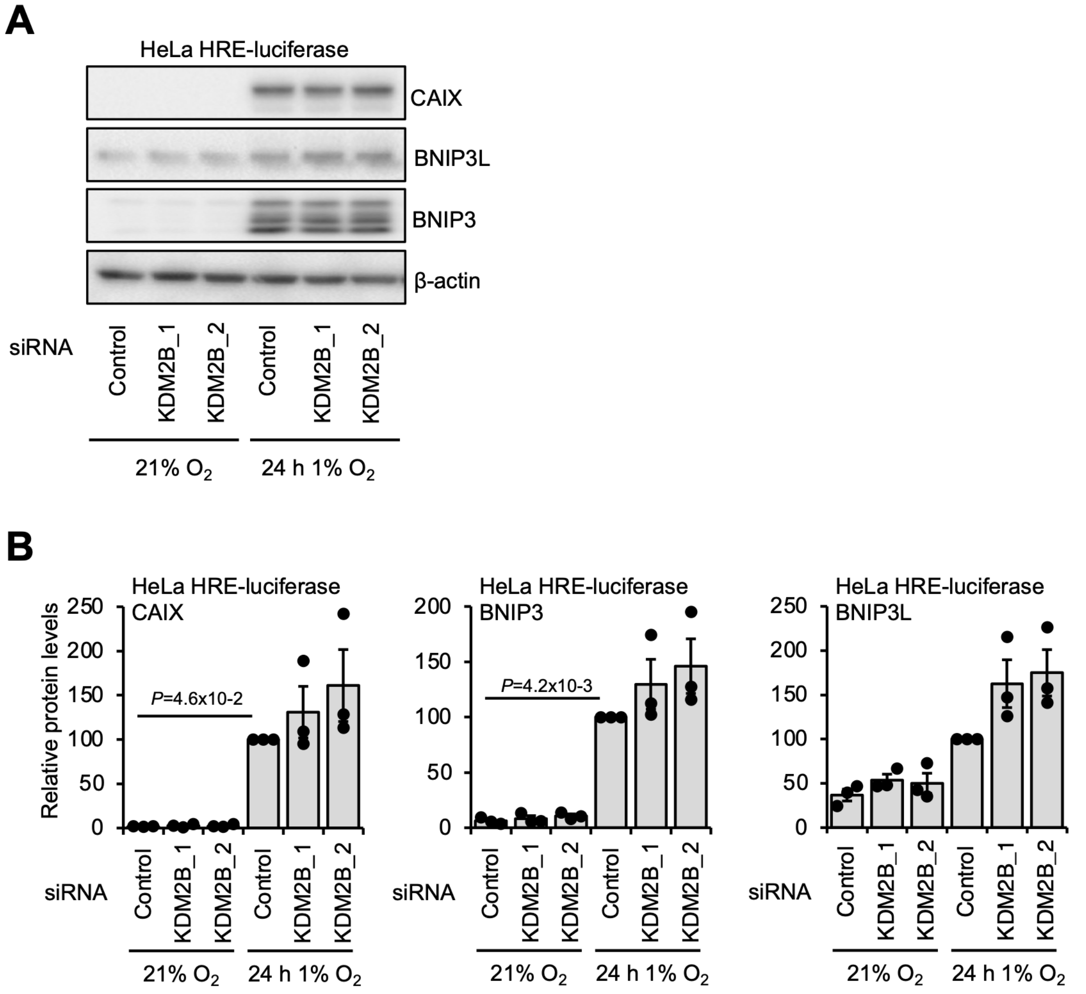


**Supplementary Figure S2. Regulation of HIF activity by KDM2B in HeLa HRE-luciferase cells**. **A)** Western blot and **B)** Western blot quantification analysis for the indicated proteins in HeLa HRE-luciferase cells cultured at 21% oxygen, transfected with control, KDM2B_1, or KDM2B_2 siRNA for 48 hours, and exposed or not to 1% oxygen for 24 hours. β-actin was used as a normalising protein. A representative Western blot of three biological replicates is shown. Data represent mean (n=3 biological replicates) ± SEM. *P* values for group comparisons were calculated by one-way ANOVA with Tukey's test.

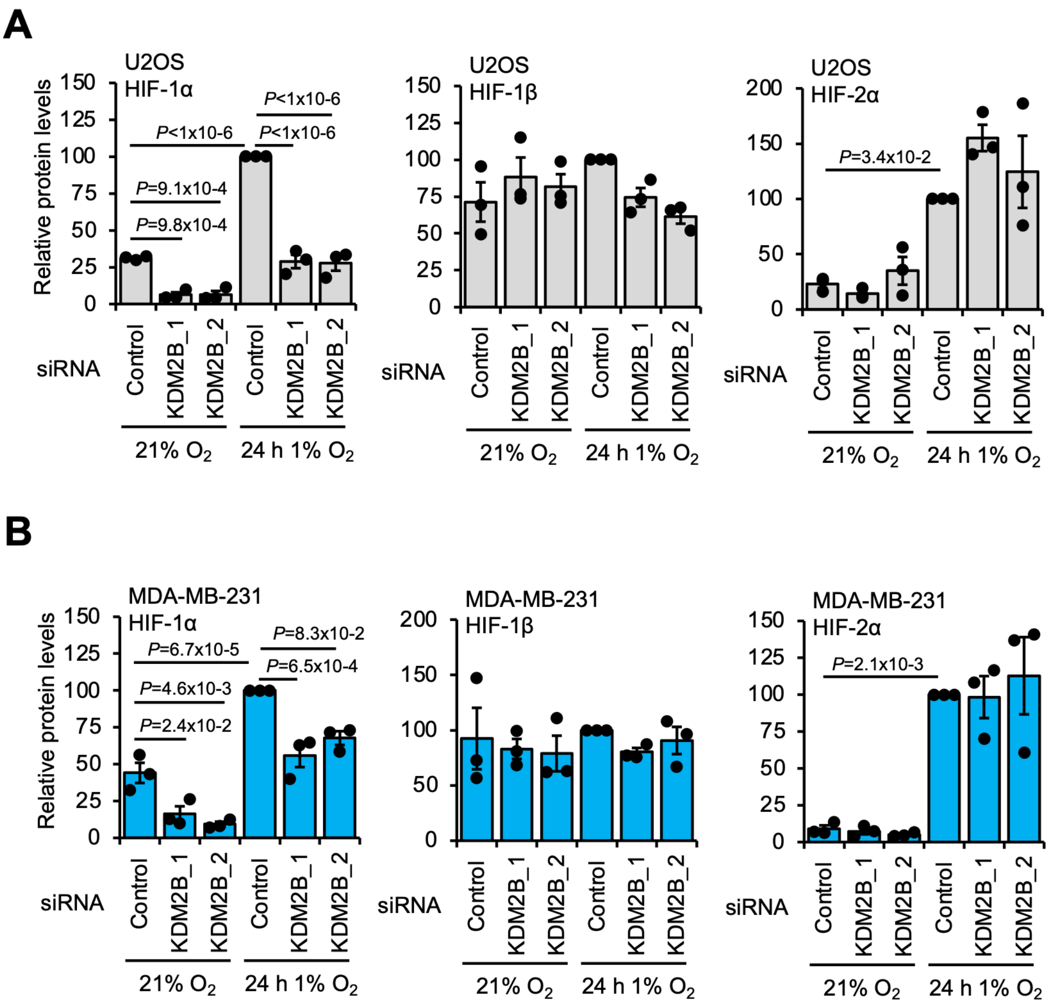


**Supplementary Figure S3. Regulation of HIF levels by KDM2B in U2OS and MDA-MB-231 cells, Western blot quantification**. Western blot quantification analysis for the indicated proteins in **A)** U2OS and **B)** MDA-MB-231 cells cultured at 21% oxygen, transfected with control, KDM2B_1, or KDM2B_2 siRNA for 48 hours, and exposed or not to 1% oxygen for 24 hours. β-actin was used as a normalising protein. Data represent mean (n=3 biological replicates) ± SEM. *P* values for group comparisons were calculated by one-way ANOVA with Tukey's test.

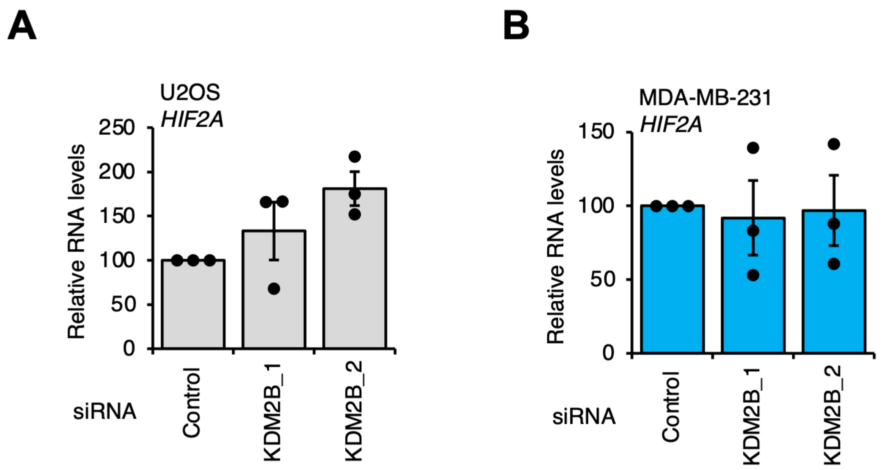


**Supplementary Figure S4. *HIF2A* RNA levels in response to KDM2B depletion.** qPCR analysis of *HIF2A* RNA levels in **A)** U2OS and **B)** MDA-MB-231 cells cultured at 21% oxygen and transfected with control, KDM2B_1, or KDM2B_2 siRNA for 48 hours. *ACTB* was used as a normalising gene. Data represent mean (*n=3* biological replicates) ± SEM. *P* values for group differences were calculated by one-way ANOVA with Dunnett's test.

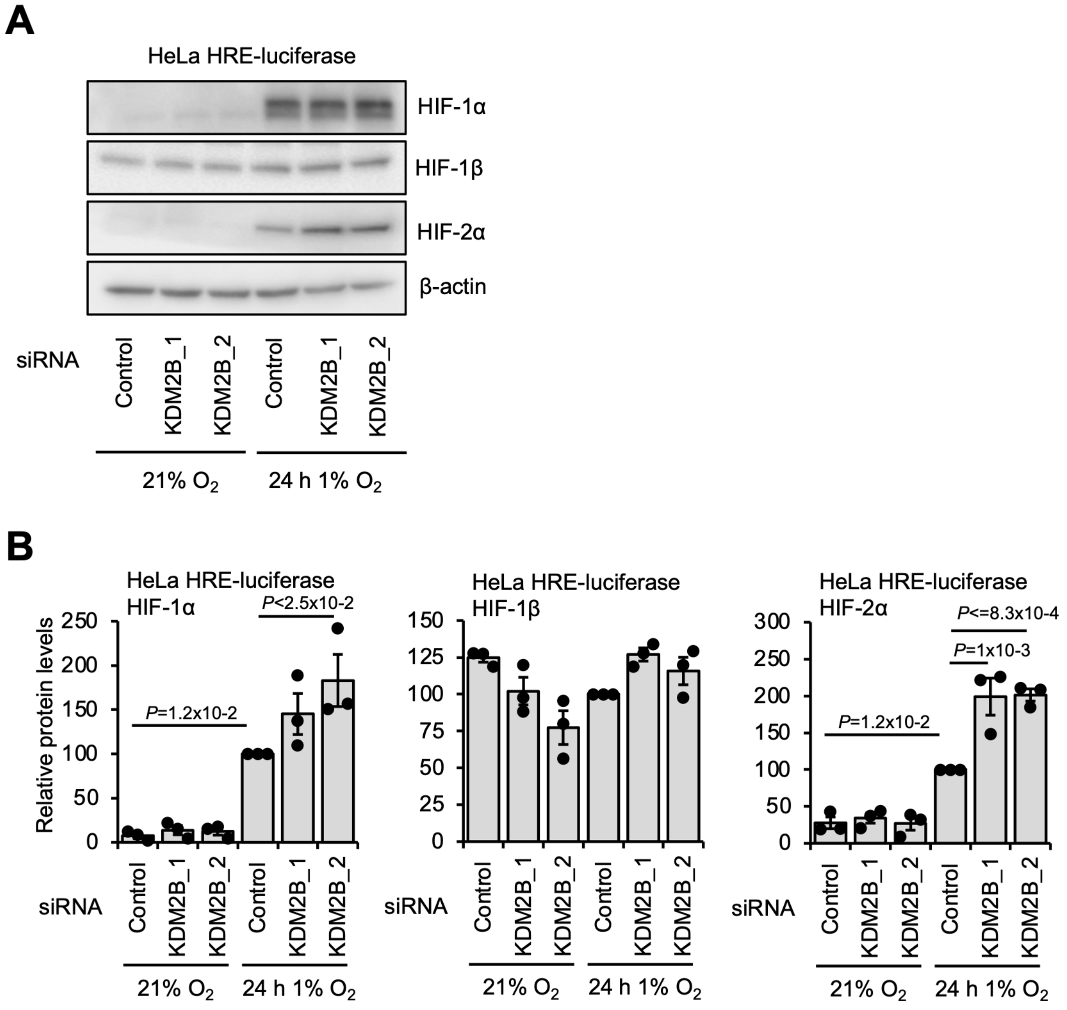


**Supplementary Figure S5. Regulation of HIF levels by KDM2B in HeLa HRE-luciferase cells**. **A)** Western blot and **B)** Western blot quantification analysis for the indicated proteins in HeLa HRE-luciferase cells cultured at 21% oxygen, transfected with control, KDM2B_1, or KDM2B_2 siRNA for 48 hours, and exposed or not to 1% oxygen for 24 hours. β-actin was used as a normalising protein. A representative Western blot of three biological replicates is shown. Data represent mean (n=3 biological replicates) ± SEM. *P* values for group comparisons were calculated by one-way ANOVA with Tukey's test.

A

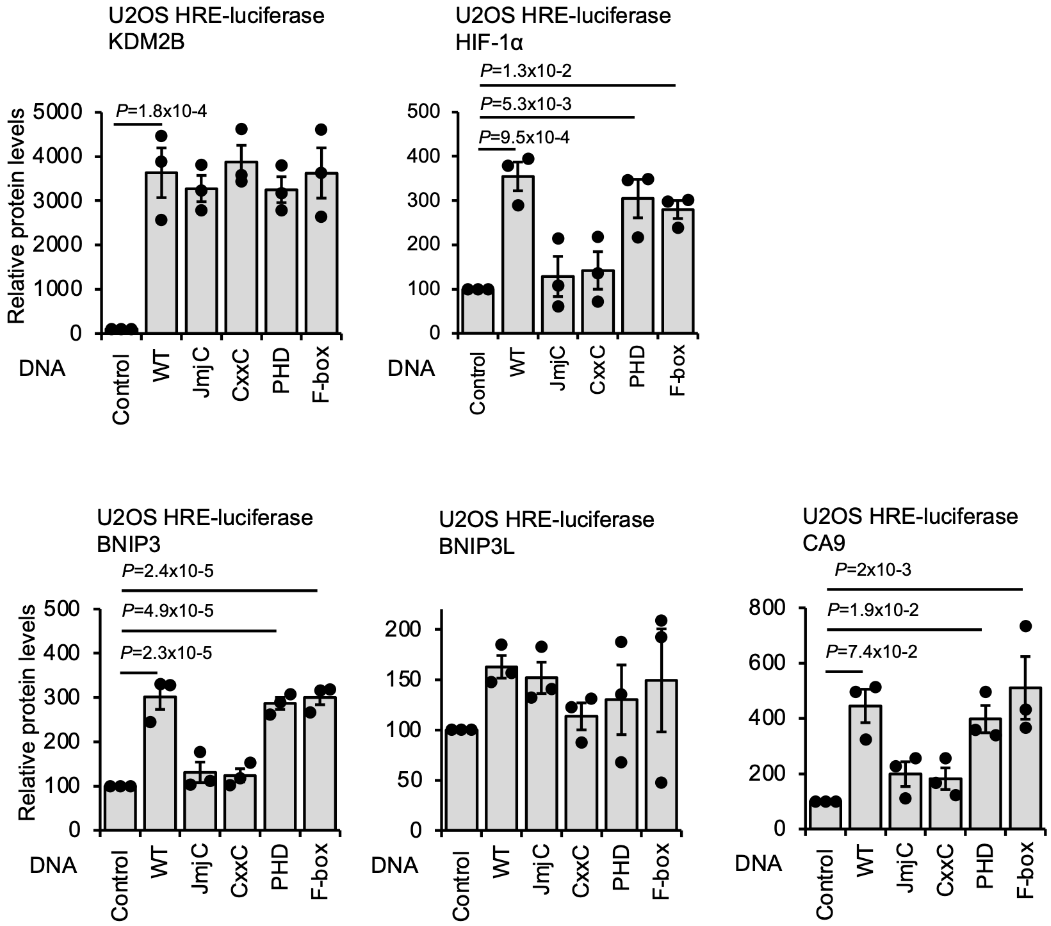


**Supplementary Figure S6. KDM2B controls HIF activity through its JmjC and CxxC domains, U2OS HRE-luciferase Western blot quantification. A)** Western blot quantification analysis for the indicated proteins in U2OS HRE-luciferase cells cultured at 21% oxygen and transfected with control, KDM2B WT, KDM2B JmjC, KDM2B CxxC, KDM2B PHD, or KDM2B F-box plasmids for 48 hours. β-actin was used as a normalising protein. Data represent mean (n=3 biological replicates) ± SEM. *P* values for group comparisons were calculated by one-way ANOVA with Dunnett's test.

**A**

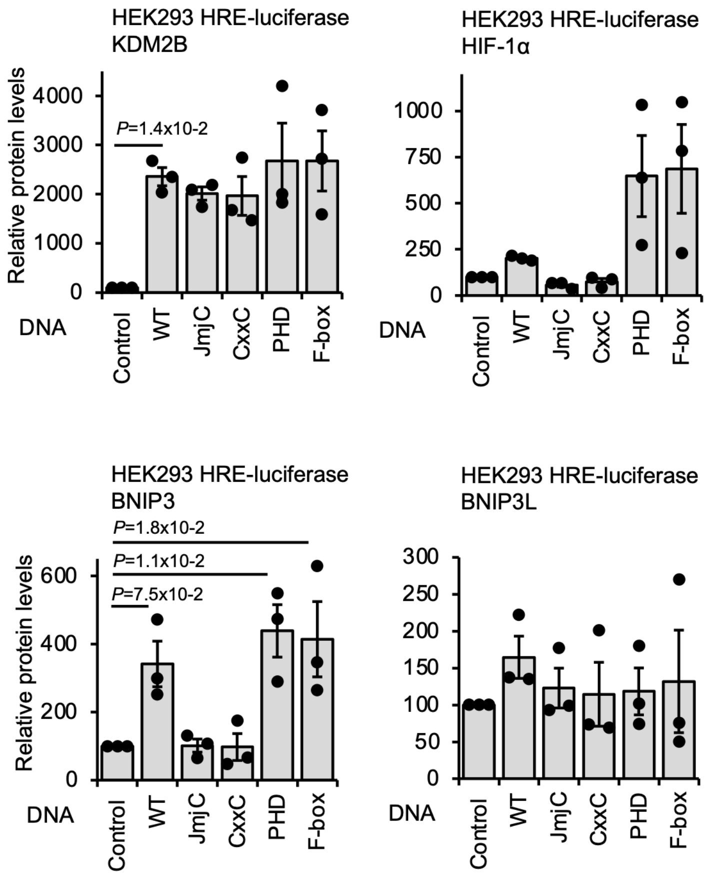


**Supplementary Figure S7. KDM2B controls HIF activity through its JmjC and CxxC domains, HEK293 HRE-luciferase Western blot quantification. A)** Western blot quantification analysis for the indicated proteins in HEK293 HRE-luciferase cells cultured at 21% oxygen and transfected with control, KDM2B WT, KDM2B JmjC, KDM2B CxxC, KDM2B PHD, or KDM2B F-box plasmids for 48 hours. β-actin was used as a normalising protein. Data represent mean (n=3 biological replicates) ± SEM. *P* values for group comparisons were calculated by one-way ANOVA with Dunnett's test.

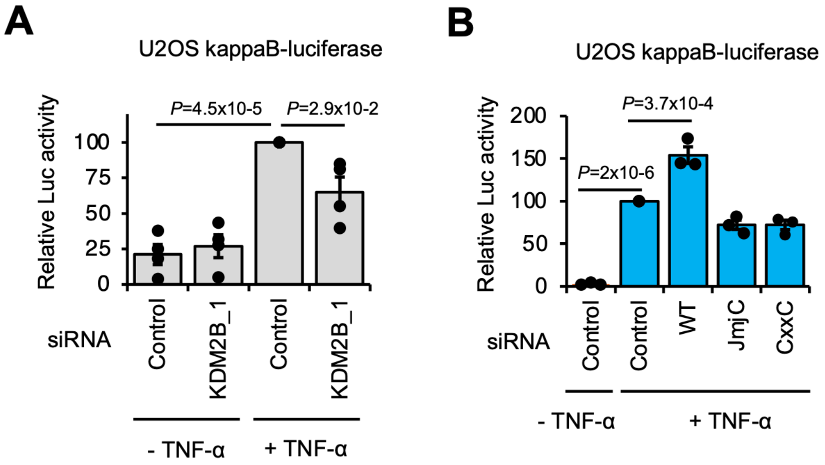


**Supplementary Figure S8. KDM2B regulates NF-κB reporter activity through its JmjC and CxxC domains. A)** Luciferase assay analysis in U2OS kappaB-luciferase cells cultured at 21% oxygen, transfected with control or KDM2B_1 siRNA for 48 hours, and treated with or without 10 ng/µL TNF-α for 8 hours. Data represent mean (n=4 biological replicates) ± SEM. *P* values for group differences were calculated by one-way ANOVA with Tukey's test. **B)** Luciferase assay analysis in U2OS kappaB-luciferase cells cultured at 21% oxygen, transfected with control, KDM2B WT, KDM2B JmjC, or KDM2B CxxC plasmids for 48 hours, and treated with or without 10 ng/µL TNF-α for 8 hours. Data represent mean (n=3 biological replicates) ± SEM. *P* values for group comparisons were calculated by one-way ANOVA with Tukey's test.

A

| Cancer Type | *R* value | *P* value |
|---|---|---|
| ACC | 0.34 | 2.2x10-3 |
| BRCA | 0.2 | 3.1x10-11 |
| CESC | 0.15 | 7.9x10-3 |
| COAD | 0.2 | 1.1x10-3 |
| DLBC | 0.6 | 1x10-5 |
| ESCA | 0.22 | 2.3x10-3 |
| HNSC | 0.13 | 2.1x10-3 |
| KICH | 0.47 | 6.5x10-5 |
| KIRC | 0.34 | 2.2x10-15 |
| KIRP | 0.28 | 1.9x10-6 |
| LAML | 0.17 | 2.5x10-2 |
| LGG | 0.09 | 4.1x10-2 |
| LIHC | 0.43 | 0 |
| LUAD | 0.11 | 1.2x10-2 |
| OV | 0.49 | 0 |
| PAAD | 0.39 | 9.8x10-8 |
| PCPG | 0.24 | 1.3x10-3 |
| PRAD | 0.33 | 1.5x10-2 |
| READ | 0.44 | 1.1x10-5 |
| SKCM | 0.25 | 9.3x10-8 |
| UCEC | 0.25 | 8.3x10-4 |
| UCS | 0.28 | 3.8x10-2 |
| UVM | 0.75 | 3.6x10-15 |

B

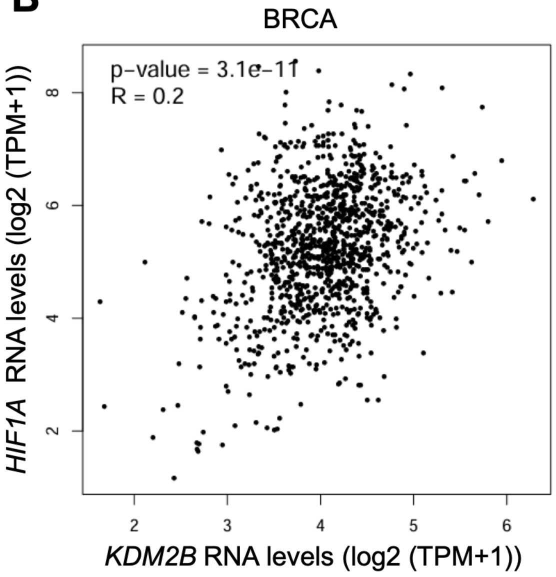


C

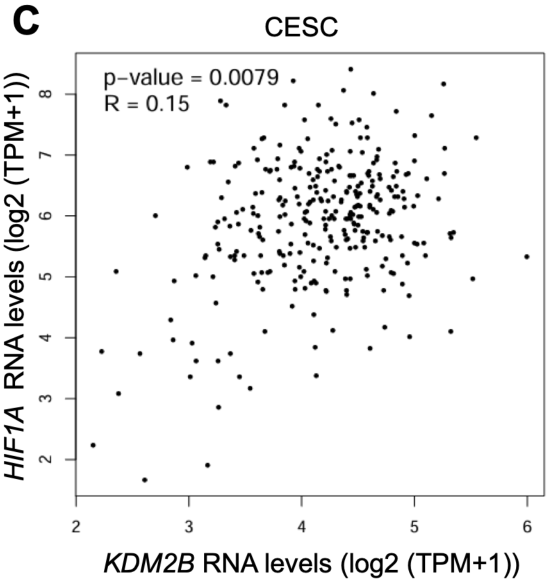


**Supplementary Figure S9. Correlation analysis between *KDM2B* and *HIF1A* RNA expression in patient cancer samples. A)** Correlation analysis between *KDM2B* and *HIF1A* RNA expression in patient samples from different cancer types from The Cancer Genome Atlas (TCGA) using GEPIA2, cancer types with a *P* value below 0.05 are shown. *P* values and *R* values were calculated using Pearson's correlation coefficient. Correlation analysis plots for **B)** BRCA (Breast Invasive Carcinoma) and **C)** CESC (Cervical Squamous Cell Carcinoma and Endocervical Adenocarcinoma). Cancer types=ACC-Adrenocortical Carcinoma; BRCA-Breast Invasive Carcinoma; CESC-Cervical Squamous Cell Carcinoma and Endocervical Adenocarcinoma; COAD-Colon Adenocarcinoma; DLBC-Lymphoid Neoplasm Diffuse Large B-cell Lymphoma; ESCA-Esophageal Carcinoma; HNSC-Head and Neck Squamous Cell Carcinoma; KICH-Kidney Chromophobe; KIRC-Kidney Renal Clear Cell carcinoma; KIRP-Kidney Renal Papillary Cell Carcinoma; LAML-Acute Myeloid Leukemia; LGG-Brain Lower Grade Glioma; LIHC-Liver Hepatocellular Carcinoma; LUAD-Lung Adenocarcinoma; OV-Ovarian Serous Cystadenocarcinoma; PAAD-Pancreatic Adenocarcinoma; PCPG-Pheochromocytoma and paraganglioma; PRAD-Prostate Adenocarcinoma; READ-Rectum Adenocarcinoma; SKCM-Skin Cutaneous Melanoma; UCEC-Uterine Corpus Endometrial Carcinoma; UCS-Uterine Carcinosarcoma; UVM-Uveal Melanoma.

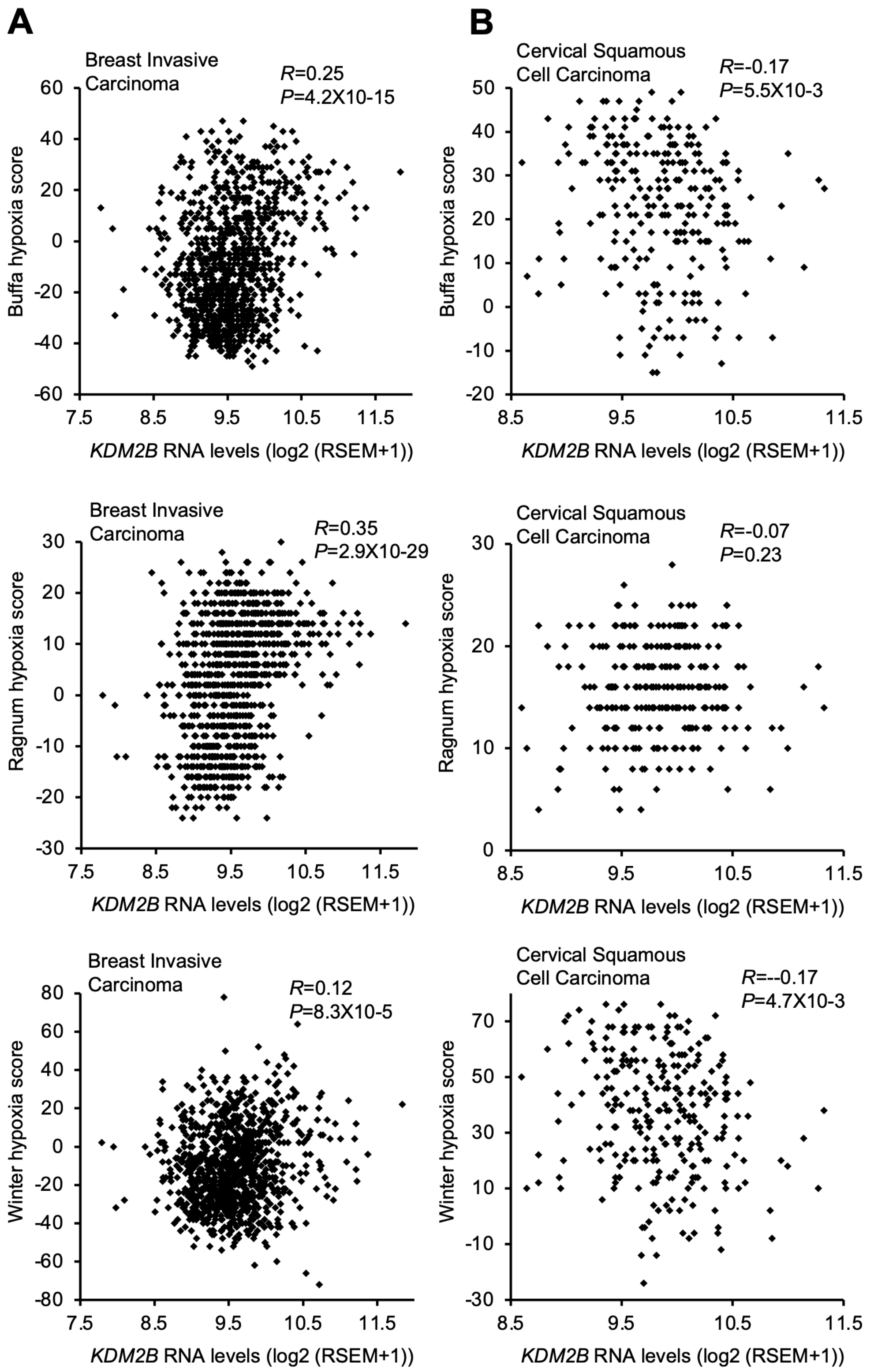


**Supplementary Figure S10. Correlation analysis between *KDM2B* and hypoxia signature RNA expression in Breast Invasive Carcinoma and Cervical Squamous Cell Carcinoma.** Correlation analysis plots between *KDM2B* and hypoxia signature (Buffa, Ragnum, and Winter) RNA expression in **A)** Breast Invasive Carcinoma and **B)** Cervical Squamous Cell Carcinoma patient samples from The Cancer Genome Atlas (TCGA) PanCancer Atlas using cBioPortal. *P* values and *R* values were calculated using Pearson's correlation coefficient.
